# Mechanical Loading Induces the Longitudinal Growth of Muscle Fibers via an mTORC1-Independent Mechanism

**DOI:** 10.1101/2025.09.19.676647

**Authors:** Jamie E. Hibbert, Kent W. Jorgenson, Ramy K.A. Sayed, Hector G. Paez, Marius Meinhold, Corey G.K. Flynn, Anthony N. Lange, Garrison T. Lindley, Philip M. Flejsierowicz, Troy A. Hornberger

**Affiliations:** Department of Comparative Biosciences, University of Wisconsin-Madison; School of Veterinary Medicine, University of Wisconsin-Madison; Department of Anatomy and Embryology, Faculty of Veterinary Medicine, Sohag University, Sohag, Egypt

## Abstract

Mechanical loading drives skeletal muscle growth, yet the mechanisms that regulate this process remain undefined. Here, we show that an increase in mechanical loading induces muscle fiber growth through two distinct mechanisms. Radial growth, reflected by an increase in fiber cross-sectional area, is mediated through an mTORC1-dependent signaling pathway, whereas longitudinal growth, marked by the in-series addition of sarcomeres, is mediated through an mTORC1-independent signaling pathway. To gain further insight into the events that drive longitudinal growth, we combined BONCAT-based labeling of newly synthesized proteins with high-resolution imaging and determined that the in-series addition of sarcomeres is mediated by a process that involves transverse splitting at the Z-lines of pre-existing sarcomeres. Collectively, our findings not only challenge the long-standing view that mechanically induced growth is uniformly governed by mTORC1, but they also lay the framework for a new understanding of the molecular and structural events that drive this process.

**Teaser:** Unlocking the Mechanical Load-Induced Growth of Skeletal Muscle: mTORC1 Doesn’t Always Hold the Key.

## Introduction

Skeletal muscle plays a vital role in numerous bodily functions such as the regulation of body temperature, breathing, voluntary movements, and metabolic homeostasis (*1, 2*). Indeed, it has been well documented that the maintenance of skeletal muscle mass is correlated with decreased morbidity and mortality risks as well as improved quality of life (*1, 3, 4*). Hence, a thorough understanding of how skeletal muscle mass is regulated has broad implications for public health.

Previous studies have established that the mechanical loads placed on the muscle are one of the most potent regulators of skeletal muscle mass, with an increase in mechanical loading leading to an increase in muscle mass and a decrease in mechanical loading leading to a decrease in mass (*5–10*). However, in many circumstances, it is not possible to directly measure changes in mass (e.g., in longitudinal studies and / or when human participants are used). Therefore, changes in whole muscle cross-sectional area (CSA) are often used as a proxy for muscle mass, and the two are often viewed as interchangeable in the literature (*11, 12*).

To date, numerous studies have been aimed at understanding how increased mechanical loading leads to an increase in muscle mass / CSA, and in most instances, it has been concluded that the increase is accompanied by a proportional increase in the amount of myofibrillar protein and / or maximal force production (*13–16*). In other words, most studies suggest that the increase in muscle mass / CSA is accompanied by a proportional expansion of the contractile machinery. In theory, this expansion could be mediated by an increase in the number of fibers in the muscle (i.e., hyperplasia), and / or an increase in the size of the fibers. However, the extent to which hyperplasia contributes to mechanically induced increases in muscle mass / CSA remains unclear (*17, 18*). For example, in one of the few studies that have painstakingly isolated and counted all of the fibers within the muscle, Gollnick et al. (1981) found that 4 weeks of chronic mechanical overload (MOV) led to a robust 45% increase in the mass of the plantaris muscle of rats, but it did not lead to a change in the number of fibers (*19*). Likewise, Timson et al. (1985) counted isolated fibers and found that 6-8 weeks of MOV in mice led to a 30-39% increase in soleus muscle mass, but again, no change in the total number of fibers was observed (*20*). Thus, while a role for hyperplasia cannot be excluded (*21*), most studies favor the conclusion that mechanically induced increases in muscle mass / CSA are primarily mediated by an increase in fiber size rather than an increase in the number of fibers.

There are two primary processes via which an increase in mechanical loading can induce an increase in muscle fiber size, and these processes are typically referred to as radial growth and longitudinal growth. Specifically, radial growth is marked by an increase in fiber CSA and involves the in-parallel addition of sarcomeric proteins, whereas longitudinal growth is marked by an increase in fiber length and involves the in-series addition of sarcomeres (*17*). Radial growth of the muscle fibers is, by far, the most widely recognized type of mechanically induced growth, and it has been observed in numerous human and animal models of increased mechanical loading (*16, 22*). However, studies in humans and animals have also provided evidence that increased mechanical loading can induce longitudinal growth and, although not intuitively obvious, the longitudinal growth of the muscle fibers can lead to substantial changes in muscle CSA (*17, 23, 24*). For example, Roy and Edgerton (1995) found that 8 weeks of MOV led to a 13% increase in the length of the fibers in the mouse plantaris muscle, and when we applied a geometric model to predict the impact that this would have on whole muscle CSA, we found that this modest increase in fiber length should lead to a 52% in mid-belly CSA (*23, 25*). Notably, the predicted increase in mid-belly CSA was not linked to any changes in fiber diameter but instead was due to a 52% increase in the number of fibers that crossed through the mid-belly of the muscle (*25*).

The aforementioned points illustrate that the response of skeletal muscle to increased mechanical loading is complex and comprised of distinct growth-related processes that work in concert. As such, it should not be surprising that the molecular mechanisms that drive this response are equally complex and remain largely undefined (*26–28*). For instance, the prevailing dogma is that mechanically induced growth will occur when the balance between protein synthesis and protein degradation is shifted in favor of a net accumulation of newly synthesized proteins (NSPs), and the mechanistic target of rapamycin complex 1 (mTORC1) is considered to be one of the most potent regulators of this balance (*5, 29, 30*). Moreover, previous studies have shown that signaling through mTORC1 is necessary for the radial growth of muscle fibers that occurs in response to MOV (*7, 31, 32*). Yet, many of the same studies have also revealed that the inhibition of signaling through mTORC1 only partially inhibits the MOV-induced increase in muscle mass, and it has no significant effect on either the MOV-induced increase in protein synthesis or the increase in the number of muscle fibers per section (a putative marker of longitudinal growth) (*7, 31*). After considering these points, we began to suspect that the longitudinal growth of fibers that occurs in response to MOV is mediated through an mTORC1-independent mechanism. Hence, the initial goal of this study was to address this possibility.

## Results

### Mechanical Overload Induces Longitudinal Growth of Muscle Fibers via an mTORC1-Independent Mechanism

To identify the growth-related adaptations that are mediated through an mTORC1-dependent pathway, mice were subjected to MOV or a sham surgery. Following the surgery, the mice were given daily injections of the mTORC1 inhibitor rapamycin or the solvent vehicle as a control condition. After 16 days, the plantaris muscles (PLT) were collected and assessed for changes in muscle mass / CSA. As shown in Figure 1A-C, the results indicated that MOV led to a robust increase in both the mass and mid-belly CSA, and these effects were only partially inhibited by rapamycin. As explained previously, the increase in muscle mass / CSA could be mediated by radial and / or longitudinal growth of the muscle fibers. Thus, to address these variables, we first measured the CSA of the primary fiber types that are found within the PLT (i.e., Type IIA, IIX, and IIB). Consistent with previous studies, the outcomes revealed that MOV induced an increase in the CSA of these fibers (i.e., radial growth), and this was effectively abolished by rapamycin (Fig. 1D) (*31, 32*). In line with earlier reports, we also determined that MOV led to a significant increase in the number of fibers per cross-section, and that this effect was mediated through a completely rapamycin-insensitive mechanism (Fig. 1E) (*31*).

**Figure 1.**
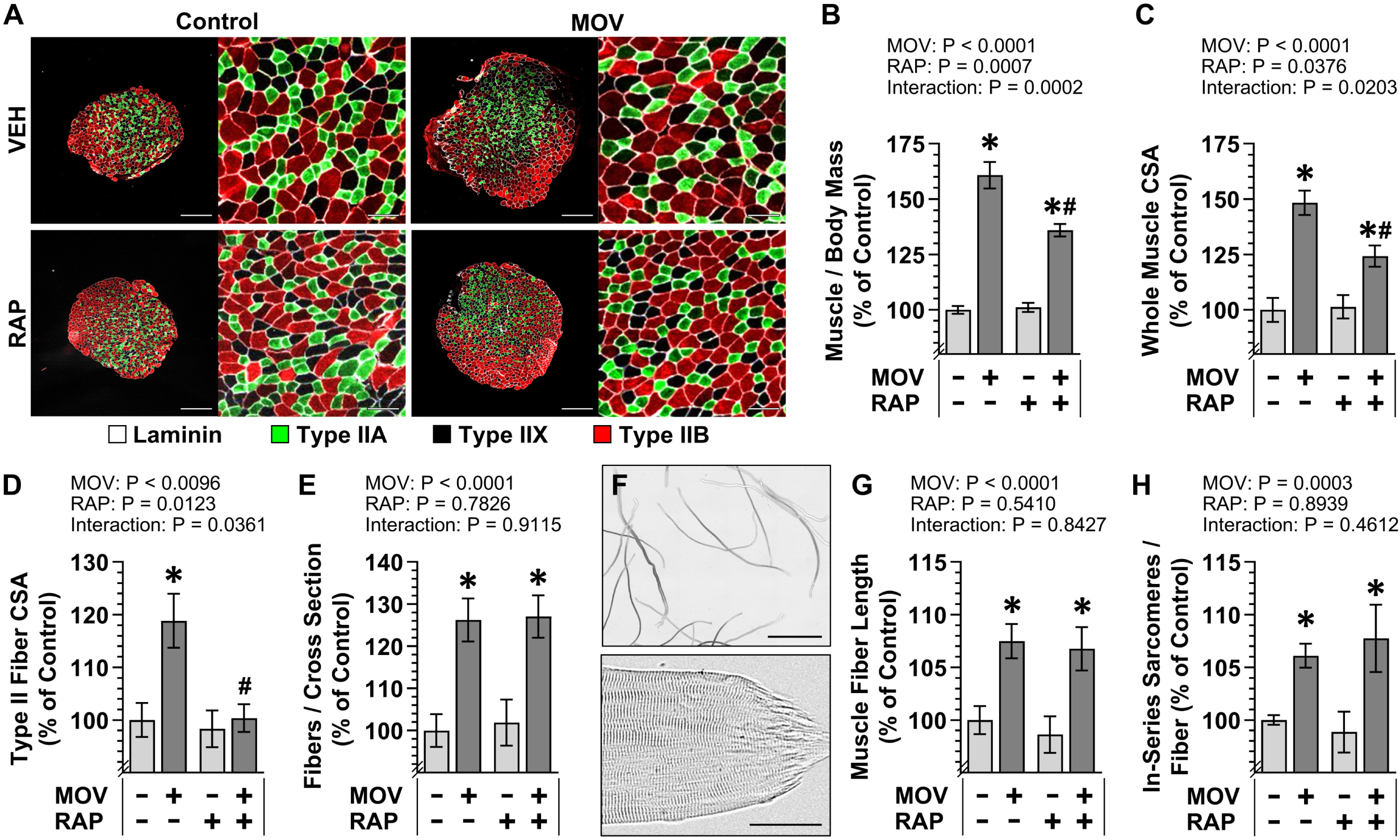
Mechanical Overload Induces Longitudinal Growth of Muscle Fibers via an mTORC1-lndependent Mechanism. C57BL/6J mice were subjected to a mechanical overload (MOV) or sham (control) surgery and injected daily with either rapamycin (RAP) or the vehicle DMSO (VEH) as a control condition. After 16 days, the plantaris (PLT) muscle from one leg was weighed, cross-sectioned at the mid-belly, and analyzed via immunohistochemistry (IHC), while the PLT from the contralateral leg was weighed and then processed for single fiber isolation and analysis via brightfield microscopy. **(A)** Representative images of cross-sections that had been subjected to IHC for laminin and fiber type identification (i.e., Type IIA, IIX, or IIB). Scale bar= 500 µm in the low magnification images and 100 µm in the high magnification images. Measurements of the **(B)** PLT muscle mass to body mass ratio, **(C)** whole muscle cross-sectional area (CSA), **(D)** the mean CSA of the Type II fibers (i.e., average of the Type IIA, IIX, and IIB fibers), and **(E)** number of fibers per whole muscle cross-section. **(F)** Representative image of isolated fibers with the higher magnification image showing an example of a “tapered end”. Scale bar= 2 mm in the low magnification image and 50 µm in the high magnification image. Measurement of the **(G)** average fiber length, and **(H)** number of in-series sarcomeres per fiber. All values are presented as group means ± SEM and expressed relative to the mean of the vehicle control condition, n = 9-36 muscles/ group. The data were analyzed with two-way ANOVA. * Significant effect of surgery within a given RAP condition,# significant effect of RAP within a given surgery condition, P ≤ 0.005.

As described in the introduction, the rapamycin-insensitive increase in the number of fibers per cross-section could potentially be explained by longitudinal growth of the muscle fibers. Thus, to directly address this possibility, we measured the length of isolated intact and full-length fibers as determined by the presence of a “tapered end” at both ends of the fibers (Fig. 1F). The results of these measurements revealed that MOV led to a 7.5% increase in the length of the fibers, and this effect was matched with a proportional increase in the number of in-series sarcomeres per fiber (Fig. 1G-H). Moreover, the outcomes revealed that the MOV-induced increase in fiber length and the increase in the number of in-series sarcomeres were mediated through a completely rapamycin-insensitive mechanism (Fig. 1F-G). Collectively, the outcomes support previous studies that have concluded that the radial growth of muscle fibers that occurs in response to MOV is mediated through a mTORC1-dependent mechanism and also reveal for the first time that the MOV-induced longitudinal growth of muscle fibers is mediated through a mTORC1-independent mechanism.

### Mechanical Overload Leads to a Progressive Increase in the Number of In-Series Sarcomeres

Having established that longitudinal growth is mediated through an mTORC1-independent mechanism, we next wanted to gain insight into the processes that drive this event. However, before we could do this, we needed to determine when the longitudinal growth was occurring. Hence, we examined fiber length, sarcomere length, and the number of in-series sarcomeres at various timepoints following the onset of MOV. As shown in Figure 2, the outcomes revealed that fiber length rapidly increased during the first 8 days of MOV. We suspected that the initial increase in fiber length might be mediated by both the in-series addition of new sarcomeres as well as an increase in sarcomere length. However, no significant differences in sarcomere length were found at any of the time-points that were examined during the 16 days of MOV (Fig. 2B). Instead, the outcomes revealed that the increase in fiber length was due to a progressive (near linear) increase in the number of in-series sarcomeres throughout the 16-day window of observation (Fig. 2C). Accordingly, we decided to focus on the middle of this window (i.e., the 8-day timepoint) for all subsequent experiments that were aimed at identifying the processes that drive longitudinal growth.

**Figure 2.**
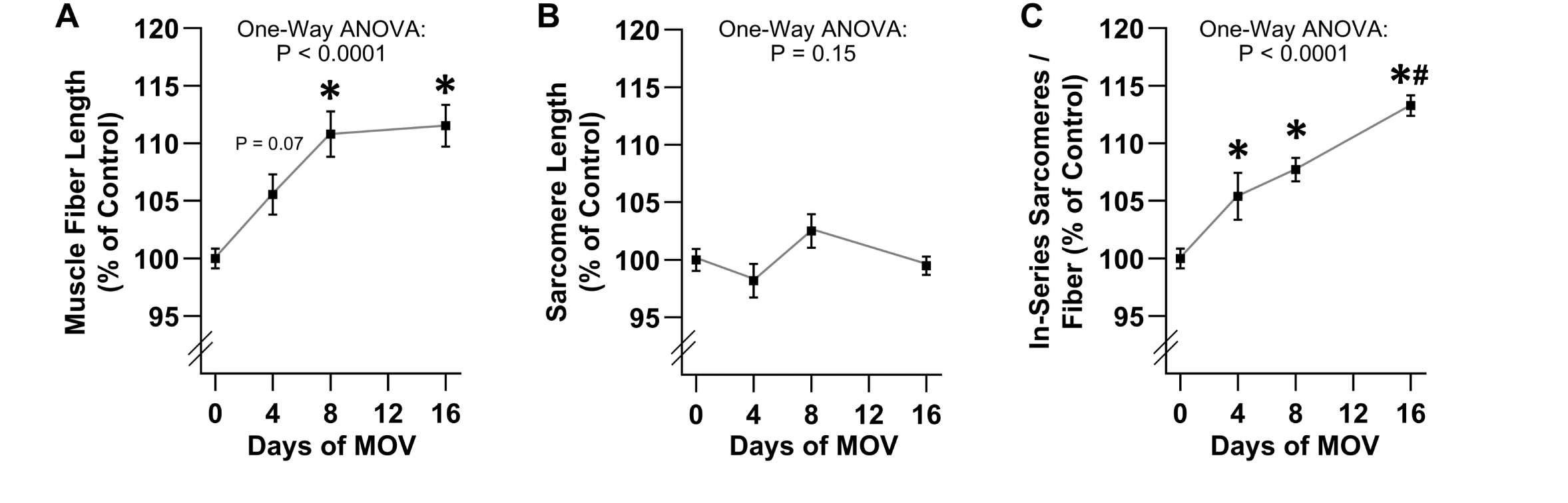
Mechanical Overload Leads to a Progressive Increase in the Number of In-Series Sarcomeres. C57BL/6J mice were subjected to a mechanical overload (MOV) or sham (control) surgery. The plantaris muscles were collected at 4, 8, or 16 days after surgery and processed for single muscle fiber isolation. Measurements of the average **(A)** muscle fiber length, **(B)** sarcomere length, and **(C)** number of in-series sarcomeres per fiber. For the sham samples, no significant difference was observed in any of the variables across the different time points. Accordingly, all of the sham samples were consolidated into a single group (i.e., 0 days of MOV). Values are presented as group means ± SEM and are expressed relative to the mean of the sham samples that were collected at each time point, n = 6-14 muscles/ group. The data were analyzed with one-way ANOVA. * Significantly different from the 0-day timepoint, # significantly different from the 8-day timepoint, P < 0.005.

### Mechanical Overload Leads to Transverse Splits / Disarray of Sarcomeres

The data in Figures 1 and 2 consistently demonstrated that MOV leads to the in-series addition of new sarcomeres. Therefore, we next wanted to gain insight into where and how the new in-series sarcomeres were being added. To date, a variety of different hypotheses have been proposed (*33–37*) and, in our opinion, the most compelling one predicts that new in-series sarcomeres are added throughout the length of the fiber via a process that involves transverse splitting of the pre-existing sarcomeres (Fig. 3A) (*17, 36–39*). Thus, to determine whether MOV led to an increase in the prevalence of transverse splits, mice were subjected to MOV or a sham surgery. After 8 days, the PLT muscles were collected, and longitudinal sections were subjected to immunohistochemistry (IHC) for α-actinin. As illustrated in Figure 3B, the resulting images were manually assessed for the net number of transverse splits that occurred across the width of the fiber and the outcomes revealed that MOV led to a 4.9-fold increase in the number of transverse splits (Fig. 3C). While conducting these analyses, we noticed that regions with extensive transverse splits could be identified by a pronounced disarray in the appearance of the Z-lines. Thus, as a complementary approach to the manual measurements, we used the ZlineDetection program developed by Morris et al. to perform automated measurements of the continuous Z-line length (*40*). Specifically, as illustrated in Figure 3D, regions with transverse splits have misaligned sarcomeres, and this misalignment / disarray leads to a reduction in the continuous Z-line length. Indeed, the same images that were used for the manual analysis of transverse splits revealed that MOV led to a 40% reduction in the average continuous Z-line length (Fig. 3E). In other words, two different analytical approaches indicated that MOV leads to a significant increase in transverse splits / disarray.

**Figure 3.**
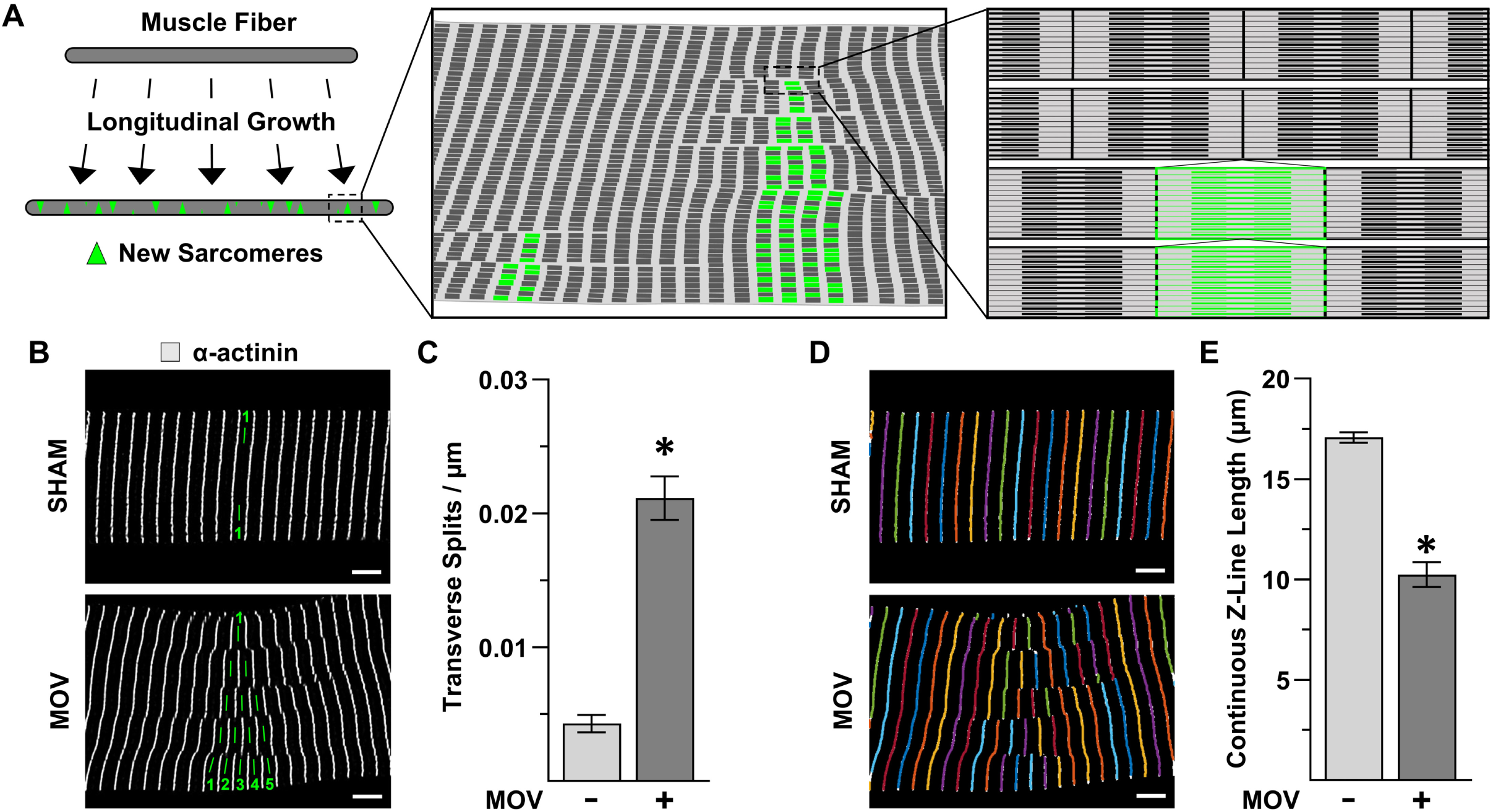
Mechanical Overload Leads to Transverse Splitting / Disarray of Sarcomeres. **(A)** Illustration of the proposed mechanism in which new in-series sarcomeres are added via a process that involves transverse splitting of the pre-existing sarcomeres. **(B-E)** C57BL/6J mice were subjected to a mechanical overload (MOV) or sham (control) surgery. At 8 days after surgery, the plantaris muscle from one leg was collected, and longitudinal sections were analyzed via immunohistochemistry (IHC). **(B)** Representative images from longitudinal sections after they had been subjected to IHC for a-actinin and subsequently had an A-band traced across the width of the fiber (note: the MOV sample shows an A-band that contains a net of four transverse splits across its width). **(C)** Manual measurements of the number of transverse splits normalized to the width of the muscle fiber. **(D)** Example of the image in B after it had been subjected to automated measurements of continuous Z-line length. **(E)** Measurements of the continuous Z-line length from the same images that were analyzed in C. All values are presented as group means ± SEM, n = 4-8 muscles/ group. The data were analyzed with student’s t-tests. * Significantly different from control, P < 0.0001. Scale bars= 5 µm.

### BONCAT-Based Approach for Visualizing and Quantifying the Accumulation of Newly Synthesized Proteins

As illustrated in Figure 3A, our hypothesis predicts that the regions with extensive transverse splits / disarray represent sites of active in-series sarcomerogenesis. To investigate this further, we needed to establish a means for identifying newly formed sarcomeres. However, this was a difficult task because no definitive markers of in-series sarcomerogenesis had previously been reported. Nevertheless, we reasoned that new sarcomeres should be largely composed of newly synthesized proteins (NSPs) and, therefore, should be overtly enriched with NSPs. Accordingly, we sought to establish a technique for visualizing and quantifying where NSPs accumulate within the muscle, and we achieved this by applying the principles of biorthogonal non-canonical amino acid tagging (BONCAT) with azidonorleucine (ANL) (*41*). Specifically, ANL is an azide-bearing analog of methionine that can be incorporated into NSPs of mice that express a mutated methionyl-tRNA synthetase (MetRS^L274G^), and the ANL in these NSPs can then be labeled using ‘click’ chemistry (Fig. 4A). To generate the mouse model, we crossed floxed STOP GFP-2A-MetRS^L274G^ mice with mice that express CMV-Cre. The presence of CMV-Cre resulted in mice with germline cells that no longer contained the STOP cassette in front of the GFP-2A-MetRS^L274G^ transgene and, in turn, enabled us to generate MetRS ^L274G+/+^ mice which ubiquitously co-express GFP and MetRS^L274G^ in the absence of Cre (Fig. 4B).

**Figure 4.**
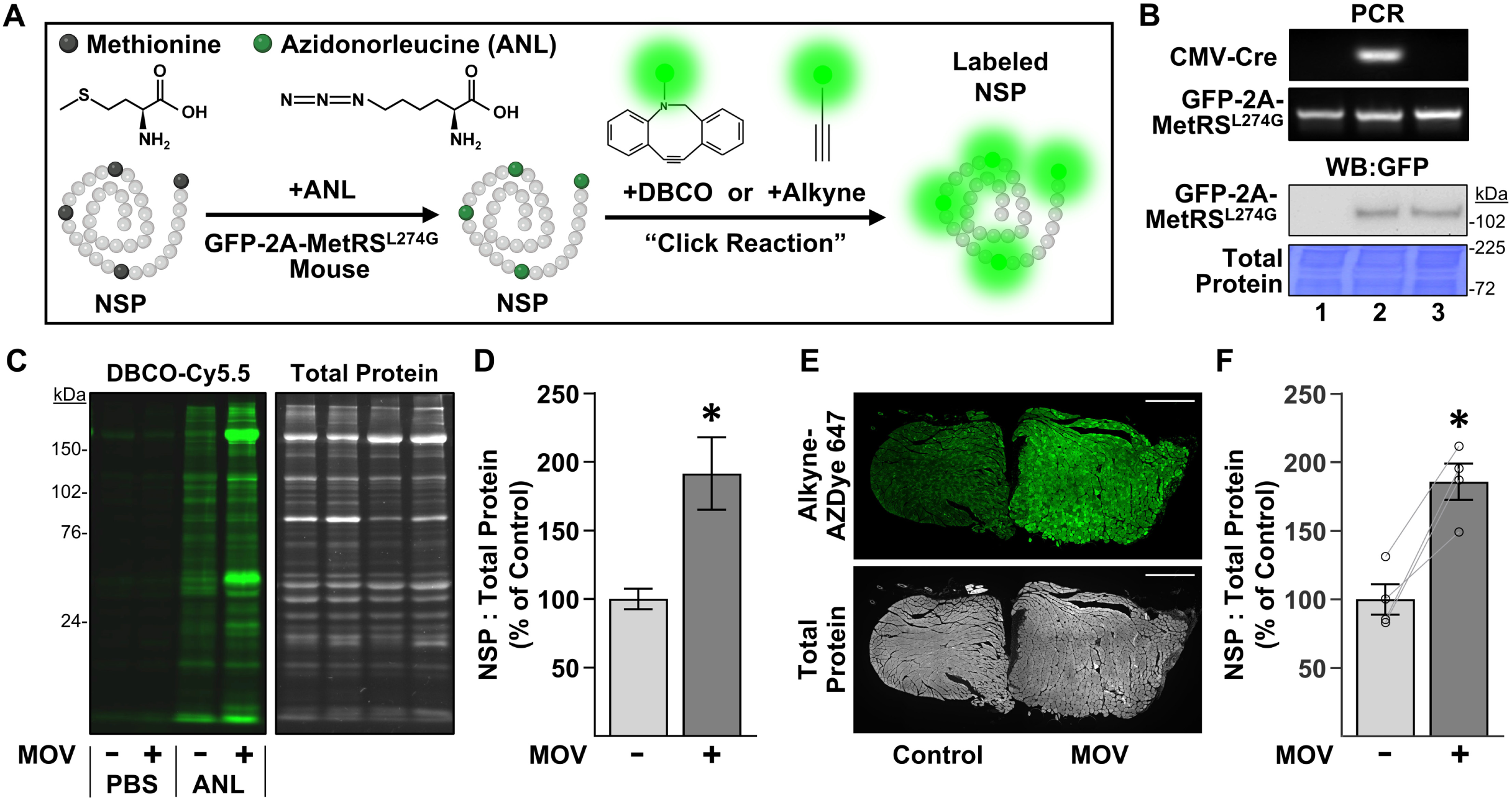
BONCAT-Based Approach for Visualizing the Accumulation of Newly Synthesized Proteins (NSPs). **(A)** Illustration of how an azide-bearing analog of methionine, called azidonorleucine (ANL), can get incorporated into NSPs in mice that express a mutated form of the methionyl-tRNA synthetase (MetRS^L274G^). Once incorporated, a “click” reaction with a fluorophore bearing a reactive DBCO or alkyne group can be used to label the azide on the ANL. **(B)** Genotyping (PCR) and western blot (WB) analysis of tails from the following: (1) floxed STOP GFP-2A-MetRS^L274G^ transgenic mice, (2) mice that express CMV-Cre plus floxed STOP GFP-2A-MetRS^L274G^, and (3) MetRS^L274G+/+^ mice that ubiquitously co-express GFP-2A-MetRS^L274G^ in the absence of CMV-Cre. **(C-F)** MetRS^L274G+/+^ mice were subjected to a mechanical overload (MOV) or sham (control) surgery. After 7 days, the mice were injected with ANL or the vehicle (PBS), and the plantaris muscles were collected 24 hr later. **(C)** Representative in-gel fluorescence images showing the DBCO-Cy5.5-labeled NSPs and total protein levels in each sample. **(D)** Quantification of the NSP to total protein ratio expressed relative to the mean of the control samples. **(E)** Representative cross-section showing Alkyne-AZDye 647-labeled NSPs and total protein from a pair of plantaris muscles (control and MOV) that were visualized with fluorescence microscopy. Scale bar = 500 µm. **(F)** The NSP to total protein ratio for each sample was expressed relative to the mean of the control samples. For illustrative purposes, the signals for Cy5.5 and AZDye 647 were pseudo colored green. Values in the bars are presented as the group means ± SEM, n = 4 muscles/ group. The data were analyzed with a student’s t-test (D) or a paired t-test (F). * Significantly different from control, P < 0.05.

To establish the utility of these mice, we subjected them to MOV or a sham surgery, and after 7 days, they were injected with ANL or the vehicle (PBS) as a negative control condition. The PLT muscles were collected 24 hours later and subjected to ‘click’ chemistry to enable the visualization and quantification of NSPs. Specifically, we used a strain-promoted ‘click’ reaction with DBCO-Cy5.5 to label the NSPs in whole muscle lysates, and then in-gel fluorescence analysis was used to visualize and quantify the results. Importantly, as shown in Figure 4C-D, the outcomes not only revealed that MOV led to a significant increase in the NSP to total protein ratio, but it also demonstrated our optimized methodology was able to detect NSPs with a high signal-to-noise ratio (i.e., the signal in ANL-injected mice compared with the PBS-injected mice). Encouraged by these results, we then sought to determine whether similar results could be obtained when NSPs were labeled in muscle cross-sections. Specifically, sham and MOV muscles were frozen adjacent to one another as pairs, and then cross-sections of the pairs of muscles were subjected to a copper-catalyzed ‘click’ reaction with Alkyne-AZDye 647 (Fig. 4E). Notably, the outcomes not only showed that MOV led to a significant increase in the NSP to total protein ratio, but they also revealed that the magnitude of this effect was very similar to the results obtained with the in-gel fluorescence analysis (Fig. 4E-F).

### Mechanical Overload Leads to the Formation of NSP Hot Spots at Sites of Transverse Splitting / Disarray

Having optimized the conditions that were needed to visualize and quantify NSPs, we returned to our question of whether the regions with extensive transverse splits / disarray were indicative of sites of active in-series sarcomerogenesis. Specifically, we subjected mice to MOV or a sham surgery, and after 7 days, they were injected with ANL. The PLT muscles were collected 24 hours later, and then longitudinal sections were subjected to IHC for α-actinin to visualize the Z-lines as well as a copper-catalyzed ‘click’ reaction to label the NSPs. Strikingly, the outcomes revealed that ⁓60% of the fibers in the MOV samples contained multiple loci that were densely populated with NSPs (i.e., NSP hot spots), whereas NSP hot spots were only present in ⁓5% of the fibers in the muscles that were subjected to the sham surgery (Fig. 5A-B).

**Figure 5.**
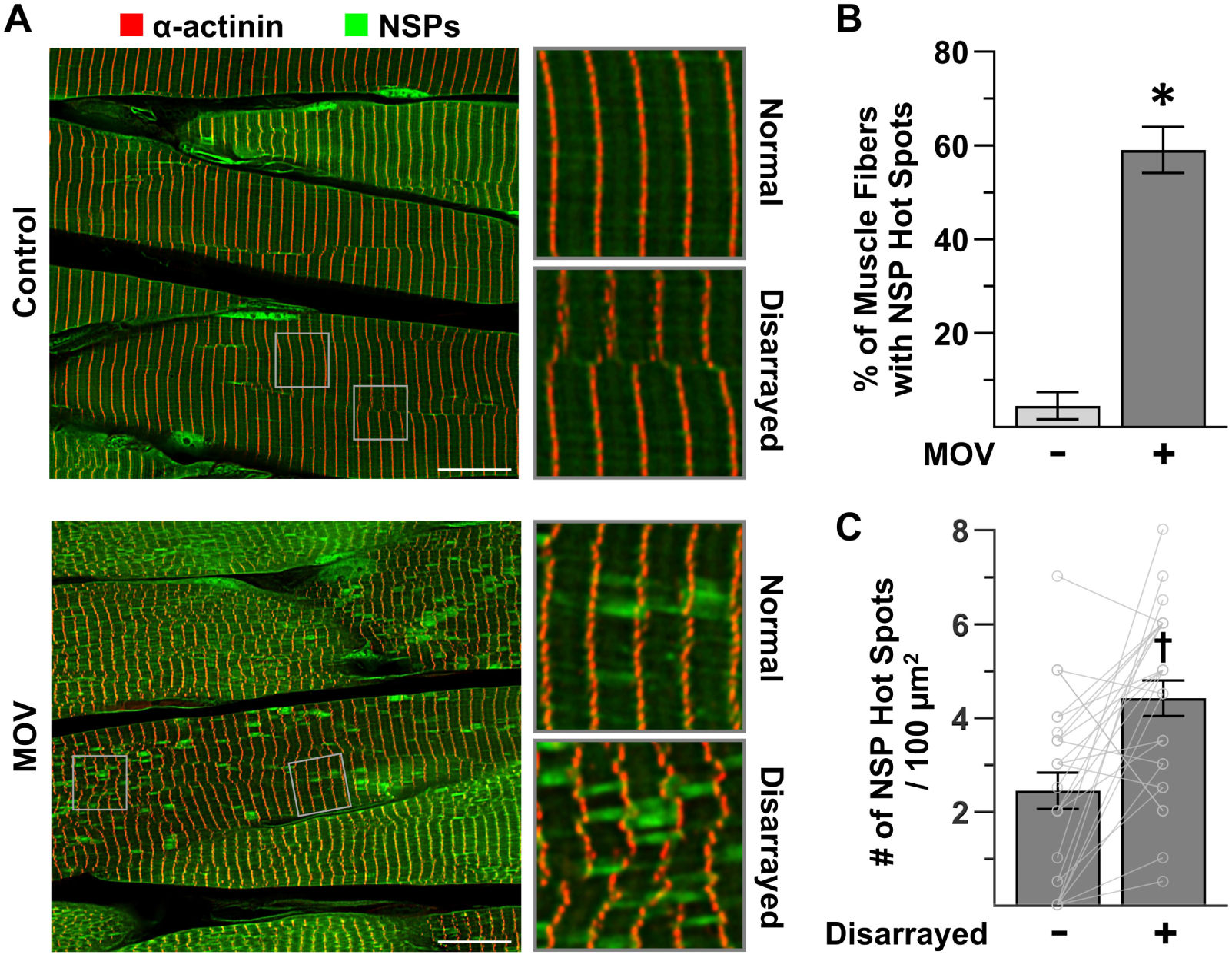
Mechanical Overload Leads to the Formation of NSP Hot Spots at Sites of Transverse Splitting / Disarray. MetRS^L274G+/+^ mice were subjected to a mechanical overload (MOV) or sham (control) surgery. After 7 days, the mice were injected with ANL, and the plantaris muscles were collected 24 hr later. **(A)** Representative images of longitudinal sections that had been subjected to immunohistochemistry for a-actinin (red) and a ‘click’ reaction with alkyne-AZDye 555 (green) to label the newly synthesized proteins (NSPs). Scale bars= 15 µm. **(B)** The percentage of fibers within each sample that were positive for NSP hot spots (n = 29-39 fibers per muscle, 3 muscles per group). **(C)** Regions of interest (ROI) within the fibers from the MOV samples that were positive for NSP hot spots were classified as having either a “normal” or a “disarrayed” Z-line configuration. An equivalent number of “normal” and “disarrayed” ROls within each fiber were then analyzed for the number of hot spots per ROI. The mean values for the “normal” and “disarrayed” ROls within each fiber were calculated, and then the mean values for each type of ROI within each fiber were reported as paired values, n = 92 ROls from 7-11 fibers per muscle, 3 muscles / group. Values in the bars are presented as the group means ± SEM, and individual paired values are shown as connected dots. The data were analyzed with a student’s t-test (B) or a paired t-test (C). * Significantly different from control, † significantly different from “normal”, P < 0.001.

According to our hypothesis, the NSP hot spots should, at least in part, be indicative of newly formed sarcomeres. By extension, this implies that the size of the NSP hot spots should resemble the size of sarcomeres (i.e., span the length between two Z-lines and have a diameter that is between ⁓400 and 1900 nm (Sup. Fig. 1)). As shown in Figure 5A, we found that the majority of the NSP hot spots were consistent with these dimensions. Our hypothesis also predicts that regions with extensive transverse splits / disarray are sites of in-series sarcomerogenesis and, as such, they should be enriched with NSP hot spots. Therefore, to test this, regions of interest (ROIs) within individual fibers from the MOV samples that were positive for NSP hot spots were classified as having either a “normal” or a “disarrayed” Z-line configuration. The number of NSP hot spots in the different types of ROIs was then quantified and, as shown in Figure 5C, the results indicated that the “disarrayed” ROIs contained significantly more NSP hot spots.

Although the above results were consistent with our hypothesis, we were surprised by the relatively high number of NSP hot spots that were observed within the ROIs that were classified as “normal”. However, our ROI classification scheme was exclusively based on the appearance of the Z-lines within the specific plane of the image that was being analyzed. Thus, we suspected that many of the observed NSP hot spots might actually be located immediately superficial or deep to a region of “disarray”. To address this, we generated 10 µm thick longitudinal sections and created three-dimensional Z-stacks that panned throughout the full depth of the sections. As shown in Supplemental Figure 2 and Supplemental Videos 1 and 2, the outcomes provided numerous examples that were consistent with our suspicion. As such, we could surmise that the magnitude of NSP hot spot enrichment that occurs at sites of “disarray” is likely much greater than what was observed with our two-dimensional quantitative approach.

### Inhibition of mTORC1 Does Not Affect the Prevalence of NSP Hot Spots or Transverse Splits in Mechanically Overloaded Muscles

Up to this point, our analyses indicated that the in-series addition of sarcomeres that occurs in response to MOV is mediated through an mTORC1-independent mechanism. Furthermore, the results provided support for our hypothesis that regions with extensive transverse splits / disarray represent sites of active in-series sarcomerogenesis. As such, we predicted that the prevalence of these regions, along with the enrichment of NSP hot spots that occurs at them, should not be altered by mTORC1 inhibition. To test this, mice were subjected to MOV or a sham surgery and received daily injections of rapamycin to inhibit signaling through mTORC1 or the vehicle (DMSO) as a control condition. After 7 days, the mice were injected with ANL, the PLT muscles were collected 24 hours later, and then mid-belly cross-sections were subjected to a copper-catalyzed ‘click’ reaction to label NSPs. As expected, the outcomes revealed that the 8 days of MOV led to an increase in the number of fibers per cross-section and that this effect was not impacted by rapamycin (Fig. 6A-B). Moreover, MOV led to an increase in both the muscle mass and the mid-belly CSA, and these effects were only partially inhibited by rapamycin (Fig. 6C-D). Notably, the lack of a full inhibitory effect of rapamycin could not be attributed to an incomplete inhibition of mTORC1, as western blot analysis confirmed that rapamycin abolished the MOV-induced increase in the phosphorylation of p70 on the threonine 389 residue - a previously validated marker of signaling through mTORC1 (Fig. 6E) (*7, 31*).

**Figure 6.**
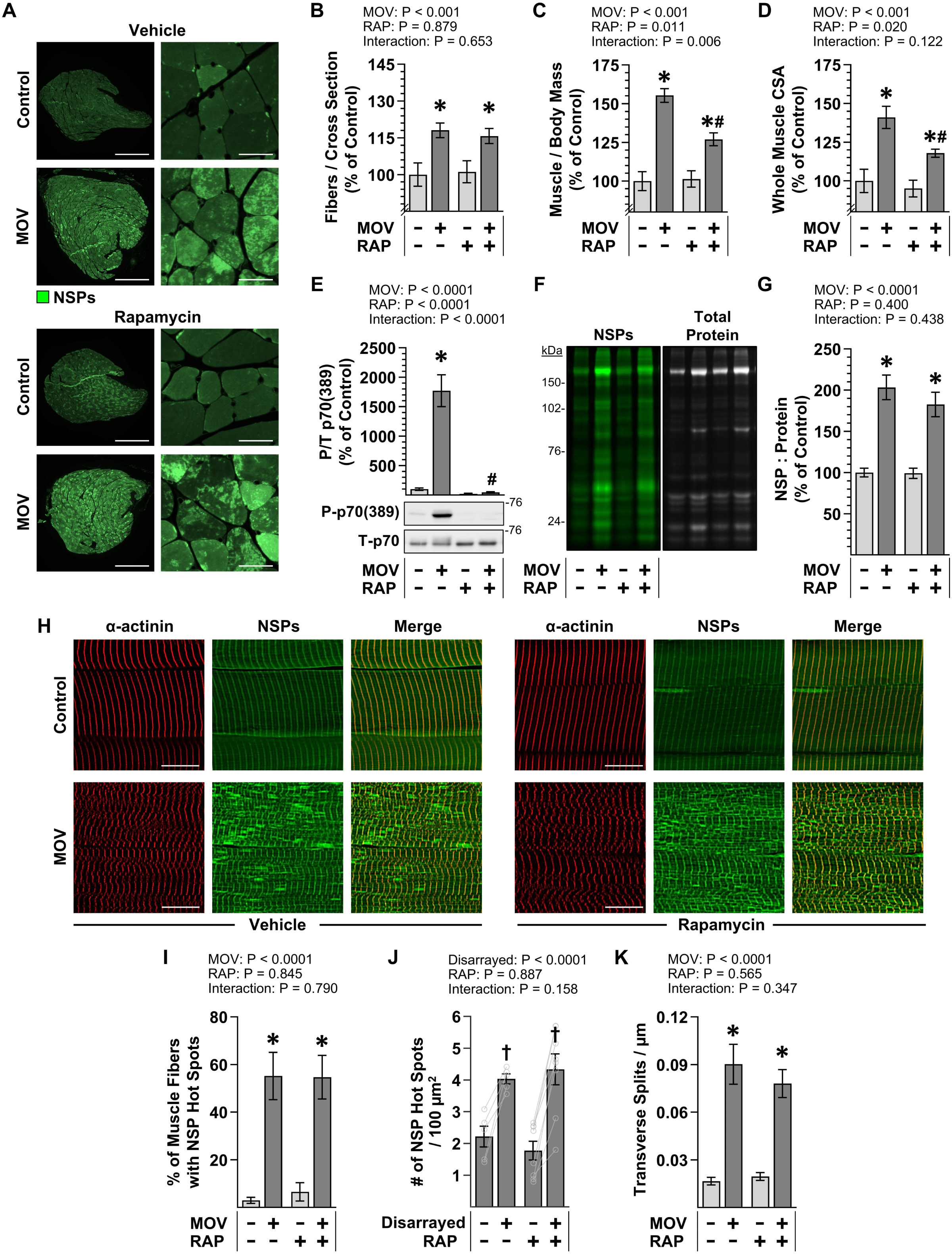
Inhibition of mTORC1 Does Not Affect the Prevalence of NSP Hot Spots or Transverse Splits in Mechanically Overloaded Muscles. MetRS^L274G+/+^ mice were subjected to a mechanical overload (MOV) or sham (control) surgery and injected daily with either rapamycin (RAP) or the vehicle DMSO (VEH) as a control condition. After 7 days, the mice were given a final dose of RAP and injected with ANL, and then the plantaris (PLT) muscles were collected 24 hr later. **(A)** Representative images of cross-sections of the PLT muscles that had been subjected to a ‘click’ reaction with Alkyne-AZDye 555 to label newly synthesized proteins (NSPs). Scale bars = 500 µm for the low magnification image and 25 µm for the high magnification images. **(B)** The number of fibers per whole muscle cross­ section, **(C)** the PLT muscle mass to body mass ratio, and **(D)** whole muscle cross-sectional area (CSA). **(E)** Representative western blots of phosphorylated (P) and total (T) levels of p70, along with the quantitative analysis of the PIT ratio. **(F)** Representative in-gel fluorescence images showing the DBCO-Cy5.5-labeled NSPs and total protein levels. **(G)** Quantitative analysis of the NSP to total protein ratio. **(H)** Representative images of longitudinal sections that had been subjected to immunohistochemistry for a-actinin (red) and a ‘click’ reaction with Alkyne-AZDye 555 (green) to label the NSPs. Scale bars = 15 µm. (I) The percentage of fibers within each sample that were positive for NSP hot spots (24-46 fibers per muscle, 6-9 muscles/ group). **(J)** 100 µm^2^ regions of interest (ROls) within the fibers from the MOV samples that were positive for NSP hot spots were classified as having a “normal” or a “disarrayed” Z-line configuration. A “normal” and a “disarrayed” ROI within each fiber was then analyzed for the number of hot spots per ROI. The mean value for the “normal” and a “disarrayed” ROls in each muscle was calculated and recorded as paired values (20-48 ROls from 10-24 fibers / muscle, 5-8 muscles / group). **(K)** The number of transverse splits normalized to the width of the muscle fiber (30 - 40 fibers/ muscle, 6-8 muscles/ group). Values were expressed relative to the mean of the vehicle control samples and presented as group means± SEM, n = 7-10 muscles/ group (B-E, G), or as the absolute group means ± SEM (I-K). Paired values are shown as connected dots (J). The data were analyzed with two­way ANOVA (B-E, G, I, and K) or two-way RM ANOVA (J). * Significant effect of surgery within a given RAP condition, # significant effect of RAP within a given surgery condition, † significantly different from “normal” within a given RAP condition, P < 0.01.

As illustrated in Figure 6A, a visual appraisal of the cross-sections suggested that rapamycin did not prevent the MOV-induced accumulation of NSPs or the formation of NSP hot spots. Thus, to formally assess this, we used a strain-promoted ‘click’ reaction to label the NSPs in whole muscle lysates and then calculated the NSP to total protein ratio in each sample. As shown in Figure 6F-G, the results indicated that MOV led to an approximate doubling in the NSP to total protein ratio, and this effect was not significantly altered by rapamycin. After obtaining this result, longitudinal sections of the PLT muscles were generated and subjected to both a copper-catalyzed ‘click’ reaction to label the NSPs and IHC to label α-actinin (Fig. 6H). Subsequent analyses of the samples demonstrated that rapamycin did not alter the MOV-induced increase in the number of NSP hot spot positive fibers, the enrichment of NSP hot spots within the regions of disarray, or the concomitant transverse splitting that leads to the regions of disarray (Fig. 6I–K). Taken together, thee results provided further support for our hypothesis that regions of transverse splits / disarray represent sites of active in-series sarcomerogenesis and demonstrate that both the structural remodeling and the localized accumulation of NSPs at these sites are mediated through an mTORC1-independent mechanism.

### The Morphology of NSP Hot Spots Reveals Potential Mechanisms of In-Series Sarcomerogenesis

At this stage, our findings consistently suggested that the NSP hot spots that accumulate at regions of transverse splits/disarray represent sites of active in-series sarcomerogenesis. To further assess the validity of this interpretation, we next evaluated how well the morphology of the NSP hot spots conformed with previously proposed models of in-series sarcomerogenesis.

The first model we considered was recently described by Rodier et al. (2025) (*42*) and is based on transverse splitting at the Z-line. Specifically, the model proposes that in-series sarcomerogenesis is initiated when titin– thick filament complexes detach from the Z-lines of an existing sarcomere. Approximately half of these complexes are pulled in one direction and the remainder in the opposite direction, thereby exposing binding sites that facilitate the recruitment of new sarcomeric proteins. If these proteins are NSPs, then the process will produce an NSP hot spot that spans the length of exactly two in-series sarcomeres (Fig. 7A).

**Figure 7.**
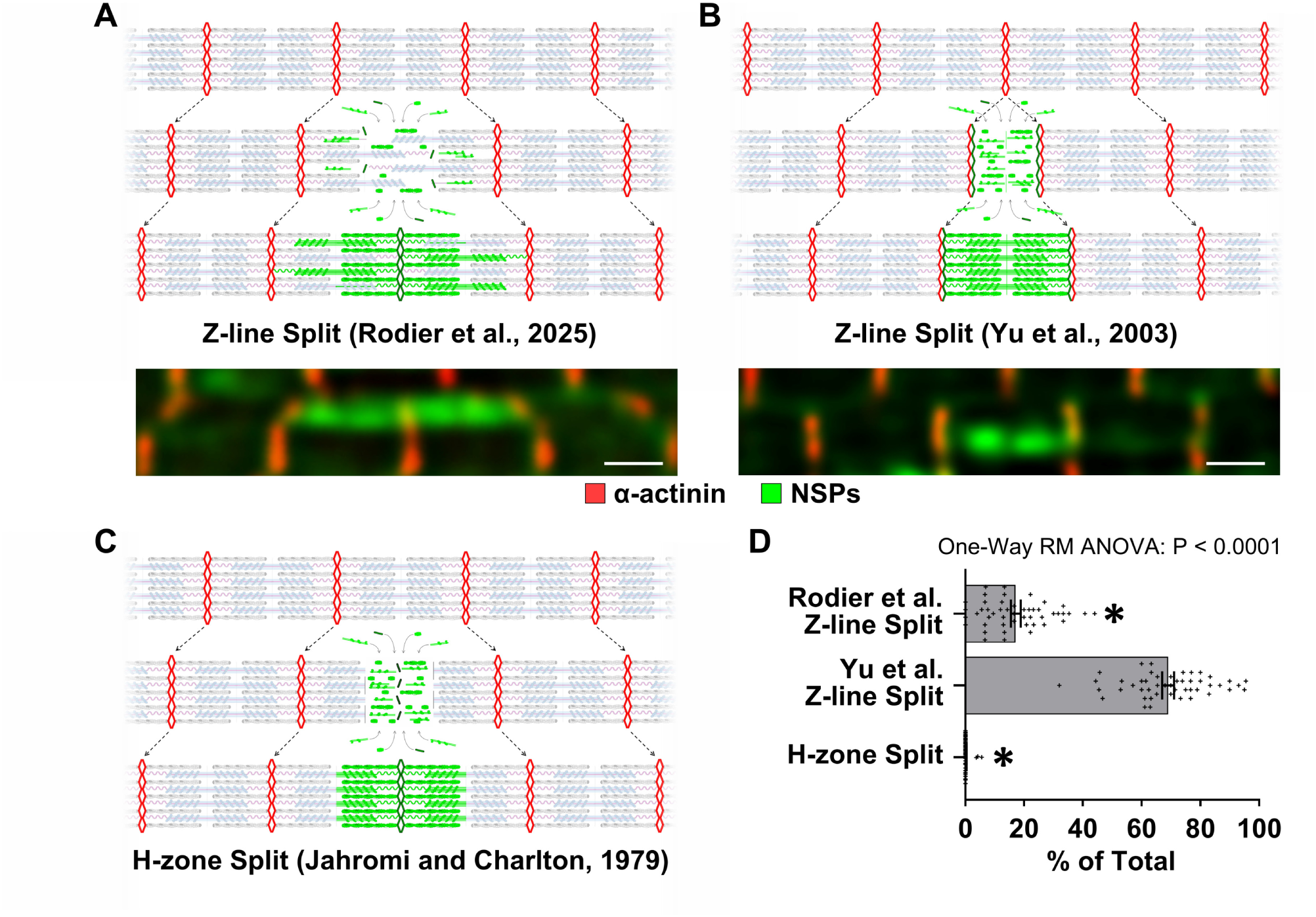
The Morphology of NSP Hot Spots Reveals Potential Mechanisms of In-Series Sarcomerogenesis. **(A-C)** Three different models of in-series sarcomerogenesis and the expected morphology of the NSP hot spots that they would produce. The schematics include Z-lines (red), titin (light pink), thick filaments (light blue), thin filaments (gray), and NSPs (green). **(A)** The Rodier et al. (2025) model of transverse Z-line splitting is expected to produce NSP hot spots that span the length of exactly two in-series sarcomeres. **(B)** The Yu et al. (2003) model of transverse Z-line splitting is expected to produce NSP hot spots that span the length of exactly one sarcomere. **(C)** The Jahromi and Charlton (1979) model of transverse H-zone splitting is expected to produce NSP hot spots that are confined to the adjacent inner halves in a pair of in-series sarcomeres. **(D)** MetRS^L274G+/+^ mice were subjected to mechanical overload (MOV), and after 7 days, the mice were injected with ANL. The plantaris muscles were collected 24 hr later, and longitudinal sections were subjected to immunohistochemistry for a-actinin (red) and a click reaction with alkyne-AZDye 555 (green) to label the NSPs. The images were assessed for the presence of fibers that had qualifying NSP hot spots (e.g., signal intensity at least 2-fold higher than the local background and dimensions that were consistent those of a sarcomere) and, for each analyzed fiber, the proportion of the NSP hot spots whose morphology aligned with each of the models of in-series sarcomerogenesis was determined, n = 48 fibers from 15-30 NSP hot spots / fiber, 10-15 fibers / muscle, 3 muscles. Representative images are shown below A and B, scale bars = 1 µm. Data are presented as individual fiber values as well as group means ± SEM. The data were analyzed with one-way RM ANOVA. * Significantly different from the Yu et al. model of Z-line splitting, P < 0.0001.

The second model, originally proposed by Yu et al. (2003) (*35*), also involves a transverse Z-line splitting event but differs in its initiation. Here, it is thought that the process begins with the breakdown and broadening of a single Z-line. As the split progresses, the residual components of the Z-line get pulled in opposite directions which, in turn, creates space for the incorporation of new sarcomeric proteins. If these proteins are NSPs, then the process will result in the formation of an NSP hot spot that spans the length of exactly one sarcomere (Fig. 7B) (*35*).

The third and final model we considered was put forward by Jahromi and Charlton (1979) (*38*) and is different from the others in that it involves transverse splitting at the H-zone. In this case, it is thought that the process begins when thick filaments get bisected at the H-zone and the resulting halves get pulled in opposite directions. During this process, new myosin molecules are incorporated at the severed ends of the thick filaments, while new thin filament–Z-line complexes get incorporated within the region that was previously occupied by the M-line. If the incorporated proteins are NSPs, then it will result in the formation of an NSP hot spot that is confined to the adjacent inner halves in a pair of in-series sarcomeres (Fig. 7C) (*38*).

After defining the morphologies of the NSP hot spots that would be expected from the different models of in-series sarcomerogenesis, we quantified the proportion of NSP hot spots in MOV muscles that conformed to each model. To this end, mice were subjected to MOV, and after 7 days, they were injected with ANL. The PLT muscles were collected 24 hours later, and then longitudinal sections were subjected to IHC for α-actinin to visualize the Z-lines as well as a copper-catalyzed ‘click’ reaction to label the NSPs. Classification analyses were then performed on NSP hot spots that exhibited a signal intensity at least two-fold above the local background and dimensions consistent with a sarcomere (400–1900 nm in diameter; see Supplemental Material 2 for details). As shown in Figure 7D, 87% of the qualifying NSP hot spots displayed morphologies that were consistent with one of the three models of in-series sarcomerogenesis. Notably, the majority of NSP hot spots aligned with the transverse Z-line splitting model proposed by Yu et al. (2003), whereas those consistent with the transverse H-zone splitting model were extremely rare (0.3%). It also bears mentioning that a subset of the NSP hot spots (13%) did not conform to any previously described model. Within this subset, approximately one-third were shorter than two sarcomeres in length and classified as “Short-Atypical,” while the remaining two-thirds extended beyond two sarcomeres and were classified as “Long-Atypical” (Sup. Fig. 3). We suspect that, rather than reflecting sites of in-series sarcomerogenesis, the atypical NSP hot spots are indicative of de novo myofibrillogenesis, a process that should culminate in long in-series streaks of newly formed sarcomeres that are densely populated with NSPs.

## Discussion

The initial goal of this study was to determine whether the longitudinal growth of muscle fibers in response to MOV is mediated through an mTORC1-independent mechanism. To investigate this, we first quantified MOV-induced changes in muscle fiber length and the number of in-series sarcomeres per fiber. As shown in Figure 1, inhibiting mTORC1 with rapamycin did not affect the MOV-induced increase in either of these markers of longitudinal growth. Notably, our previously described geometric model of the plantaris predicted that the seemingly modest MOV-induced increase in fiber length (7.5%) would result in a 27% increase in the number of fibers per cross-section (*17, 25*). Consistent with this prediction, the outcomes of our studies revealed that MOV led to a 26% increase in the number of fibers per cross-section and that the inhibition of mTORC1 did not alter this effect. In stark contrast, the inhibition of mTORC1 abolished the radial growth of the fibers that occurred in response to MOV. This was a significant distinction because it not only clarified why the inhibition of mTORC1 only partially inhibited the MOV-induced increase in muscle mass and whole muscle CSA, but, more importantly, it provided vital support for our conclusion that the longitudinal and radial growth of muscle fibers are regulated by distinct signaling pathways.

Once we had established that the longitudinal growth that occurs in response to MOV is mediated through an mTORC1-independent mechanism, we then wanted to gain insight into the mechanisms that drive this event. During this pursuit, we found that MOV leads to an increase in transverse splits, and this results in the formation of regions that are marked by misalignment of the Z-lines (i.e., disarray). We also determined that the inhibition of mTORC1 did not alter the prevalence of the regions with extensive transverse splits / disarray or the enrichment of NSP hot spots that occurs at them. Importantly, regions of disarray have been observed in other studies following mechanical overload and have often been labeled as sites of damage (*43–45*). However, this is not always the case. For instance, using electron microscopy, Yu et al. (2004) found regions of disarray in human skeletal muscles several days after a bout of intense resistance exercise, and concluded that rather than being sites of damage, these regions were indicative of adaptive myofibril remodeling (*37*). Thus, while we acknowledge that MOV can cause damage, the findings presented throughout our study have led us to conclude that the enrichment of NSP hot spots at regions of disarray is primarily indicative of active in-series sarcomerogenesis, rather than damage-induced remodeling.

To further evaluate the validity of our position, we examined whether the morphological features of the NSP hot spots aligned with previously described models of in-series sarcomerogenesis (*35, 38, 42*). As shown in Figure 7, we found that 87% of the analyzed hot spots displayed morphologies that matched the expectations of these models. This finding not only strengthened our position but it also provided insight into the mechanism that likely drives in-series sarcomerogenesis. Specifically, nearly 70% of the hot spots exhibited morphologies that were consistent with the expectations of the transverse Z-line splitting model originally proposed by Yu et al. (2003) (*35*). The high frequency of this morphology alone suggests that a mechanism that mirrors the basic features of the Yu et al. model is the predominant driver of MOV-induced in-series sarcomerogenesis. In further support of this idea, we found that only 0.3% of the hot spots had a morphology that aligned with the expectations of transverse H-zone splitting, indicating that this mechanism plays little, if any, role. Instead, the next most common morphology (17%) aligned with the expectations of the Rodier et al. (2025) model of transverse Z-line splitting. However, it is important to note that two adjacent Yu et al. Z-line splits would produce the same morphology as a Rodier et al. Z-line split. Thus, a significant proportion of the NSP hot spots that were classified as Rodier et al. Z-line splits may have actually been the result of adjacent Yu et al. Z-line splits.

It is also worth emphasizing that the Rodier et al. model, while highly compelling, was derived from observations that were made during the developmental growth of indirect flight muscles in *Drosophila* (*42*). This is important to consider because fundamental differences in mammalian and invertebrate flight muscles make the occurrence of a Rodier et al. Z-line split in mammals less likely. For example, the Rodier et al. model proposes that Z-line splits begin with a rupture in the weakest tension-bearing linkages in the sarcomere, and the interface that exists between myosin and the titin homolog Sallimus was identified as one of the most likely sites of rupture (*42*). However, in mammalian skeletal muscle, such a rupture is less likely because, unlike Sallimus which only attaches to the ends of thick filaments, mammalian titin binds to myosin along the entire length of the thick filaments. Furthermore, indirect flight muscles express a short and stiff isoform of Sallimus that lacks the spring-like PEVK domain (*46*). Consequently, the Sallimus–myosin linkage, with fewer contact points and more rigid components, should be more susceptible to rupture than the elastic titin–myosin interaction that is present in mammals (*46*). Thus, when taken together, our findings consistently suggest that the MOV-induced enrichment of NSP hot spots at regions of disarray is due to in-series sarcomerogenesis, and that the new sarcomeres are formed via an mTORC1-independent mechanism that predominantly mirrors the features of the transverse Z-line splitting model originally described by Yu et al. (2003) (*35*).

In addition to providing insight into the mechanisms that drive in-series sarcomerogenesis, our study also lays the foundation for a potentially transformative view of the role that mTORC1 plays in mechanically induced growth. This is especially important because the prevailing dogma holds that such growth will arise when the balance between protein synthesis and protein degradation shifts toward a net increase in protein synthesis, and for the last two decades, numerous studies have identified mTORC1 as a potent regulator of this balance. For instance, the activation of mTORC1 can promote an increase in protein synthesis via enhanced translation initiation and elongation, as well as through the induction of ribosome biogenesis (*47–49*). Simultaneously, the activation of mTORC1 can suppress protein degradation by inhibiting autophagy and restraining the ubiquitin– proteasome system (*50*). Given these effects, along with previous studies which have shown that the inhibition of mTORC1 prevents the radial growth of fibers that occurs in response to MOV, it is not surprising that mTORC1 has been widely viewed as a key regulator of mechanically induced growth (*29, 51, 52*). Unexpectedly, however, several recent studies have shown that various forms of mechanical stimuli (e.g., resistance exercise, MOV, and even endurance exercise) can promote an increase in protein synthesis via a fully mTORC1-independent mechanism (*7, 53–56*). Consistent with these studies, our results demonstrate that the enhanced accumulation of NSPs that occurs at 7–8 days after the onset of MOV is unaffected by mTORC1 inhibition (Fig. 6). Since the induction of longitudinal growth was also unaffected by mTORC1 inhibition, it made sense that the accumulation of the NSPs that contribute to this type of growth remained intact. What remains unclear, however, is how mTORC1 inhibition can prevent radial growth without a concomitant reduction in the accumulation of NSPs. Currently, the mechanism responsible for this remains a mystery, but a potential explanation could reside within the ability of mTORC1 to suppress protein degradation. Specifically, under normal conditions, MOV robustly activates signaling through mTORC1 (Fig. 6), and this, in turn, would be expected to suppress the activation of protein degradation. However, if signaling through mTORC1 is blocked (e.g., by rapamycin), then the suppressive effect of mTORC1 would be lost, and this should lead to an excessively large MOV-induced increase in protein degradation. As illustrated in Supplemental Figure 4, we have envisioned how such an effect would preferentially block the radial growth of muscle fibers and, if correct, this would indicate that the distinct roles that mTORC1 plays in the radial and longitudinal growth of muscle fibers is not related to its effects on protein synthesis, but instead, is uniquely dependent on its ability to suppress protein degradation. Moving forward, it will be important to test this hypothesis as the outcomes could reshape our understanding of the role that mTORC1 plays in the mechanically induced growth of skeletal muscle.

While the results of the experiments presented here are novel, they are not without limitations. For instance, our conclusions about the mechanism of MOV-induced in-series sarcomerogenesis were inferred from the morphologies of NSP hot spots; however, we cannot definitively prove that the NSP hot spots mark newly formed sarcomeres. Unfortunately, no alternative markers of newly formed sarcomeres are available to further test our conclusions, and as such, the identification of new markers of sarcomerogenesis and / or the development of new technologies for visualizing sarcomerogenesis will be needed to further test the validity of our conclusions. It is also important to consider that our primary conclusions were derived from observations that were made with the MOV model. Since the MOV model applies chronic mechanical overload to the muscle, the outcomes observed with this model may not reflect what occurs during other forms of mechanically induced growth (e.g., resistance exercise, reloading following disuse, etc.) (*57*). However, this concern is partially mitigated by the fact that our observations strongly aligned with a Yu et al. model of transverse Z-line splitting, a model that was derived from observations made in human skeletal muscles that had been subjected to a bout of intense resistance exercise (*35, 37, 39, 45*). Nonetheless, further studies will be needed to determine whether the mechanisms identified in our study are conserved across other models of increased mechanical loading.

In summary, our study demonstrates that MOV induces muscle fiber growth through two distinct mechanisms. Radial growth, reflected by an increase in fiber CSA, is mediated through an mTORC1-dependent mechanism, whereas longitudinal growth, marked by the in-series addition of sarcomeres, is mediated through an mTORC1-independent mechanism. Moreover, by integrating high-resolution imaging and BONCAT-based labeling of NSPs, we have gained new insight into the mechanisms that regulate the different forms of growth. Together, our findings have not only challenged the long-standing view that mechanically induced growth is uniformly governed by mTORC1, but they have also laid the foundation for a new understanding of the molecular and structural events that drive this process.

## Materials and Methods

### Animals

All animal experiments followed the guidelines approved by the Institutional Animal Care and Use Committee of the University of Wisconsin-Madison (#V005375). Wild type male C57BL/6J mice were obtained from Jackson Laboratory (strain #000664). MetRS^L274G^ ^+/+^ mice were generated through a multi-step breeding approach in which floxed STOP GFP-2A-MetRS^L274G^ mice (Jackson Laboratory, strain #028071) (*58*) were initially crossed with mice that express CMV-Cre (Jackson Laboratory, strain #006054). The presence of CMV-Cre resulted in offspring with germline cells that contained the recombined variant of the allele (i.e., it no longer contained the STOP cassette in front of the floxed STOP GFP-2A-MetRS^L274G^ transgene). The mice with germline transmission of the recombined allele (i.e., GFP-2A-MetRS^L274G^) were then crossed to create offspring that were homozygous for GFP-2A-MetRS^L274G^ (MetRS^L274G+/+^). Only male MetRS^L274G+/+^ offspring or male wild type C57BL/6J were used for the experiments in this study. Genotypes were confirmed with tail snips followed by PCR using the primers listed in Supplemental Material 1. All mice were housed under controlled conditions (25°C, 12 hr light/dark cycle with lights off at 6:00 PM) and provided standard rodent chow (Inotiv Teklad 2020x) and water *ad libitum*. All mice within a given intervention were randomly assigned to an experimental group and when needed, euthanasia was performed with cervical dislocation on mice that were anesthetized with isoflurane or ketamine plus xylazine as detailed below.

### Mechanical overload

Bilateral mechanical overload (MOV) surgeries were performed between 8:00 AM and 12 PM as previously described (*59*). Specifically, mice were anesthetized with 1-5% isoflurane mixed with oxygen, and then the plantaris muscles (PLT) were subjected to MOV by removing the distal one-third of the gastrocnemius muscle while leaving the soleus and plantaris muscles intact. Mice in the control groups were subjected to a bilateral sham surgery where an incision was made on the lower leg and then closed. Following the surgeries, incisions were closed with non-absorbable nylon sutures (Oasis) and Vetbond Tissue Adhesive (3M). After 4-16 days of recovery, the mice were anesthetized with 1-5% isoflurane mixed with oxygen, and then the PLT muscles were collected and processed, as described below. As with the surgeries, all terminal collections were performed between 8:00 AM and 12 PM.

### Rapamycin injections

Rapamycin (L.C. Laboratories) was dissolved in DMSO to generate a 5 μg/μl stock solution. For all relevant experiments, 1.5 mg/kg of rapamycin in 200 μl of PBS (or an equivalent amount of DMSO in 200 μl PBS for the vehicle control condition) was administered via an intraperitoneal (IP) injection immediately before the mice were subjected to mechanical overload or the sham surgery (*60*). These injections were repeated every 24 hr for up to 16 days.

### Muscle fiber type-specific analyses

Upon collection, the PLT muscles were immediately submerged in optimal cutting temperature (OCT) compound (Tissue-Tek; Sakura) at resting length and then frozen in liquid nitrogen-chilled isopentane. Mid-belly cross-sections (10 μm thick) were taken perpendicular to the long axis of the muscle with a cryostat and immediately fixed in −20°C acetone for 10 min. Immunohistochemistry for individual fiber types and subsequent analyses were performed, as previously described (*7*). Specifically, images of the entire cross-section were captured, and then the area of the entire muscle cross-section, along with the total number of muscle fibers per cross-section, were determined. The cross-sectional area (CSA) of at least 70 randomly selected fibers of each fiber type (Type IIA, IIX, or IIB) was also measured by tracing the periphery of the fiber with Nikon NIS-Elements D software. All image analyses were performed by investigators who were blinded to the experimental conditions of the samples.

### Single muscle fiber isolation and analysis

Isolated single fibers from the PLT muscle were obtained with modifications of the method described by Wada et al. (2002) (*61*). Specifically, the skin on the hindlimbs was retracted, and the hindlimb was severed at the hip and fixed in solution A (4% paraformaldehyde, 137 mM NaCl, 5.4 mM KCl, 5 mM MgCl2, 4 mM EGTA, 5 mM HEPES, pH 7.2) for 3 hr at room temperature on an orbital shaker at 150 rpm. After fixation, the limb was washed three times in phosphate-buffered saline (PBS) for 1 min each, and then the PLT muscle was removed, and measurements of mass and length were recorded. The muscle was then digested for 3 hr at room temperature in a microcentrifuge tube that contained 1 mL of 40% NaOH and placed on an orbital rocker at 150 rpm. After digestion, the muscle was washed with 1 mL of 20% NaOH for 10 min with a gentle inversion of the tube once per minute. The solution was then removed, and the muscle was transferred to a new microcentrifuge tube containing 1 mL of PBS. The PBS was then very carefully removed and replaced with 1 mL of fresh PBS. The tube was then placed on a vertical rotator and allowed to rotate for at least 12 hr at 4°C. Once the muscle had completely dissociated, a wide p200 tip (a regular p200 pipette tip with the distal 15 mm cut off) was used to transfer 50 µl of the solution to a glass slide, which was then very gently overlayed with a 22 x 22 mm cover slip, and the excess solution was wicked away with the edge of a Kimwipe. The cover glass was then sealed with clear nail polish, and an automated Keyence BZ-X microscope was used to collect and combine 20x images from the entirety of the region encased with the cover slip. The resulting images were manually searched for intact and full-length fibers as determined by the presence of a “tapered end” at both ends of the fibers (Fig. 1F), and then the length of such fibers was measured (⁓30 fibers per muscle). The mean sarcomere length for each of these fibers was also determined by recording the average sarcomere length along three linear segments within each fiber. The mean sarcomere length was then used to determine the total number of in-series sarcomeres per fiber (i.e., the fiber length divided by the mean sarcomere length). All image analyses were performed by investigators who were blinded to the experimental conditions of the samples.

### Immersion fixation and cryoprotection

Immersion fixation of hindlimbs was performed as we have previously described (*59*). Briefly, each hindlimb was collected by retracting the skin and then severing at the hip. The foot and proximal end of the femur were then secured onto a circular piece of aluminum wire mesh (Saint-Gobain ADFORS insect screen) with 90° angles between the foot-tibia and tibia-femur joints using 4-0 non-absorbable silk sutures (Fine Science Tools). The hindlimb was then submerged in a glass beaker filled with 20 mL of a solution containing 4% paraformaldehyde in 0.1 M phosphate buffer (PB) (25 mM sodium phosphate monobasic, 75 mM sodium phosphate dibasic, pH 7.2) for 3 hr at room temperature on an orbital shaker at 50 rpm. The PLT muscle was then extracted from the hindlimb and submerged in a microcentrifuge tube containing 1 mL of 4% paraformaldehyde in 0.1 M PB. The tube was then placed on a nutating rocker for 21 hr at 4°C. The PLT muscle was subsequently cryoprotected by placing it in a microcentrifuge tube containing 1 mL of 15% sucrose in 0.1 M PB. The tube was placed on a nutating rocker for 6 hr at 4°C, and then the solution was switched to 1 mL of 45% sucrose in 0.1 M PB. The tube was rocked at 4°C for an additional 18 hr, and then the muscle was rapidly immersed in OCT (Tissue-Tek), frozen in liquid nitrogen-chilled isopentane, and stored at -80°C, as previously described (*62*).

### Perfusion fixation and cryoprotection

This procedure involved slight modifications of the workflow that we recently detailed in Zhu et al. (2025) Specifically, silicone tubing was connected to a peristaltic pump with one end of the tubing connected to a 23-gauge needle (Sol-Millennium Medical Inc., product # 110201030032) while the other end was inserted into a flask containing 30 mL per mouse of ice-cold solution B (PBS, 25 mM β-glycerophosphate, 25 mM NaF, 1 mM Na_3_VO_4_) and the tubing was cleared of air bubbles. Each mouse was then weighed and anesthetized with an IP injection of a mixture containing 300 mg/kg ketamine and 60 mg/kg xylazine. Complete anesthesia was verified by checking corneal and pedal pain reflexes and, if needed, further anesthesia was induced with an additional IP injection of 50 mg/kg ketamine or via 2% isoflurane in oxygen. At this point, the PLT from one leg was quickly removed, immediately frozen in liquid nitrogen, and stored for subsequent homogenization. A tourniquet was then tightly secured around the thigh of the leg from which the PLT had been removed, and then the mouse was placed in a supine position, and their contralateral hindlimb was secured with 90° angles between the foot-tibia and tibia-femur joints. Next, an incision was made into the abdominal cavity, followed by cutting through the diaphragm and ribcage to expose the heart. The needle was then inserted into the apex of the left ventricle, the right atrium was cut, and perfusion was initiated at 7 mL/min and continued with all 30 mL of solution B. The perfusate was then switched to 30 mL of ice-cold solution C (4% paraformaldehyde in 0.1 M PB (pH 7.2) plus 25 mM β-glycerophosphate, 25 mM NaF, 1 mM Na_3_VO_4_). After the perfusion was complete, the hindlimb was skinned, excised from the mouse, and fixed in solution C for 3 hr at 4°C with orbital shaking at 50 rpm. The PLT was then removed and incubated overnight on a nutating rocker at 4°C in a microcentrifuge tube that contained 1 mL of solution C. After this incubation period, the PLT was cryoprotected, frozen in OCT, and stored at -80°C as described in the preceding section.

### Transverse splitting

Longitudinal sections (3 µm thick) from the central portion of the cryoprotected PLT muscles were collected on microscope slides with a cryostat that was chilled to -35°C. Immediately upon collection, the sections were transferred to diH_2_O to hydrate the samples. While ensuring that the sections remained hydrated, the area on the slide surrounding the section was dried with a Kimwipe, and then a hydrophobic circle was drawn around the section using an Aqua Hold 2 pen (Scientific Device Laboratory). The slides were then placed into a humidified box for all subsequent washing and incubation steps, which were performed at room temperature on a rotating rocker set to 50 rpm. The sections were incubated for 30 min in solution D (0.5% Triton X-100 and 0.5% bovine serum albumin (BSA) dissolved in PBS) and then overnight in solution D containing anti-α-actinin mouse IgG1 (1:150, Novus #NBP1-22630). The next day, the sections were washed 3 x 5 min and then 3 x 1 hr and subsequently incubated overnight in solution D containing Alexa 488 Conjugated Goat Anti-Mouse IgG1 (1:2000, Jackson Immunoresearch #115-545-205). The next day, the sections were washed 3 x 5 min and then 3 x 1 hr and subsequently incubated for 20 min with PBS containing Phalloidin-CF680R (1:50, Biotium #00048). Finally, after 2 x 5 min washes with solution D and 2 x 5 min washes with PBS, the sections were mounted with Prolong Diamond (ThermoFisher) and a 12mm coverslip and allowed to cure for 48 hr. The signals for α-actinin and Phalloidin in randomly selected 63x fields were then imaged with a Leica THUNDER Imager Tissue 3D microscope. Z-stacks with a step size of 0.22 µm were captured through the entire thickness of the sample. Then, deconvolution was applied to the Z-stack with Leica’s Small Volume Computational Clearing (SVCC) algorithm (refractive index set to 1.47, strength set to 80-100%, and regularization set to 0.05). For each sample, a total of 40 randomly selected A-bands from 4 independent fields were identified, and the average net number of transverse splits that occurred across the width of the fiber was determined, as illustrated in Figure 3. Importantly, all image acquisitions and analyses were performed by investigators who were blinded to the experimental conditions of the samples.

### Continuous Z-line length

The same 63x images that were previously analyzed for transverse splitting were also evaluated for continuous Z-Line length using the ZlineDetection program developed by Morris et al. (*40*), and downloaded from Github (https://github.com/Cardiovascular-Modeling-Laboratory/ZlineDetection) as the “zlineDetection-master” ZIP file. Specifically, for each image, a 200 x 400-pixel (i.e., 20.6 x 41.2 µm) region within each fiber was randomly selected by a blinded investigator and oriented so that the Z-lines ran perpendicular to the long axis of the selection (9-35 fibers per muscle). The image for each fiber was then analyzed according to the instructions in the userGuide.pdf provided in the ZIP file, and the average continuous Z-Line length from all of the analyzed fibers in each sample was calculated.

### Western blot analysis

Tails were homogenized with a Polytron for 20 sec in chilled solution E (8 M Urea, 50 mM Tris (pH 8.0), 25 mM β-glycerophosphate, 25 mM NaF, 1 mM Na_3_VO_4_, 10 µg/ml leupeptin, and 1 mM PMSF), centrifuged at 6,000g for 2 min, and placed on a nutating rocker for 90 min at 4°C for complete dissolution. Skeletal muscles were homogenized with a Polytron for 20 sec in ice-cold solution F (40 mM Tris (pH 7.5), 1 mM EDTA, 5 mM EGTA, 0.5% Triton X-100, 25 mM β-glycerophosphate, 25 mM NaF, 1 mM Na_3_VO_4_, 10 µg/ml leupeptin, and 1 mM PMSF). The concentration of proteins in the resulting homogenates was determined with a DC protein assay kit (Bio-Rad, Hercules, CA, USA). Equivalent amounts of protein from each sample were then dissolved in Laemmli buffer, heated to 100°C for 5 min, and subjected to electrophoretic separation by SDS-PAGE. Following electrophoretic separation, proteins were transferred to a PVDF membrane, blocked with 5% powdered milk in Tris-buffered saline containing 0.1% Tween 20 (TBST) for 1 hr followed by an overnight incubation at 4°C with one of the following primary antibodies dissolved in TBST containing 1% BSA ((anti-GFP (1:1000, Cell Signaling #2555), anti-total p70 (1:2000, Cell Signaling #2708), or anti-phospho p70(389) (1:1000, Cell Signaling #9234)). The membranes were then washed for 30 min in TBST and probed with a peroxidase-conjugated secondary antibody (1:5000, Vector Laboratories #PI-1000) for 1 hr at room temperature. Following 30 min of washing in TBST, the blots were developed with an iBright™ FL1500 system or a Chemi410 camera mounted to a UVP Autochemi system (UVP, Upland, CA, USA) using regular enhanced chemiluminescence (ECL) reagent (Pierce, Rockford, IL, USA) or ECL Prime reagent (Amersham, Piscataway, NJ, USA). Once the appropriate image was captured, the membranes were stained with Coomassie Blue to verify equal loading in all lanes. Images were quantified using ImageJ software (U.S. National Institutes of Health, Bethesda, MD; http://rsb.info.nih.gov/nih-image/).

### Azidonorleucine (ANL) injection

ANL (Iris Biotech #HAA1625) was dissolved in PBS to generate a 300 mM stock solution. For all relevant experiments, 400 µg/g bodyweight in 200 μl of PBS (or an equivalent amount of PBS for the vehicle control condition) was administered via an IP injection 24 hr before collection.

### In-gel quantification of newly synthesized proteins with BONCAT

The analyses were performed with slight modifications of our previously described protocol (*60*). Briefly, muscles were homogenized in ice-cold solution F and assessed for the concentration of proteins, as described above. An aliquot of each sample containing 134 µg of protein was then precipitated with ice-cold acetone (4:1 volume to volume ratio), centrifuged at 12,000g for 5 min, and then the acetone was removed. The pellet was air-dried for 5 min and subsequently resuspended in 90 µL of freshly prepared solution G (8 M urea, 50 mM Tris (pH 8), 100 mM 2-chloroacetamide, 25 mM NaF, 15 mM β-glycerophosphate, 10 µg/ml leupeptin, 1 mM Na_3_VO_4_, and 1 mM PMSF). Once the pellet had dissolved, the ANL-labeled proteins were labeled with Cy5.5 by adding 10 µL of a DBCO stock solution (134 µM Cy5.5 DBCO (Click Chemistry Tools) dissolved in PBS containing 40% DMSO). The combined solution was protected from light and incubated on a vertical rotator (9 rpm) for 1 hr at room temperature. Following the incubation, the proteins were precipitated with 900 µL of 100% methanol and then centrifuged at 12,000g for 5 min. The supernatant was removed, and the pellet was washed by resuspending it in 900 µL of 100% methanol and centrifuging at 12,000g for 5 min. The supernatant was again removed, and the pellet was air-dried for 5 min and then resuspended in 100 µL of solution H (40 mM Tris (pH 7.5), 1 mM EDTA, 5 mM EGTA, 0.5% Triton X-100, and 2% SDS). The resulting sample was dissolved in Laemmli buffer, boiled for 5 min, and subjected to SDS-PAGE on a separating gel that contained 8% acrylamide in the upper half and 12% acrylamide in the lower half. The gel was run at 100 V until the dye front was 5 mm from the bottom of the gel. No-Stain™ (Invitrogen) was then used, according to manufacturer instructions, to label all proteins in the gel. The gel was imaged with the universal mode of an iBright™ FL1500 (Invitrogen) and the default filters for a No-Stain labeled gel, as well as Alexa Fluor 680 (610-660 nm excitation, 710-730 nm emission) for the visualization of the Cy5.5 (i.e., the ANL-labeled proteins). Images were quantified using ImageJ software.

### Histological labeling of newly synthesized proteins with BONCAT

Cross-sections and/or longitudinal sections (3 µm thick) from the central portion of cryoprotected PLT muscles were collected on microscope slides with a cryostat that was chilled to -35°C. Immediately upon collection, the sections were transferred to diH_2_O. Residual water around the fully hydrated section was removed, and a hydrophobic circle was drawn around each section using a PAP pen (Thermo Fisher Scientific). The sections were washed with PBS for 5 min and blocked for 30 min at room temperature in solution D. The sections were then washed with PBS (3 x 5 min) and subjected to a copper-catalyzed ‘click’ reaction to tag the ANL-labeled proteins. Specifically, a 300 μL reaction mixture (enough for 6 sections) was prepared by adding the following reagents in the order listed along with vortexing between each addition: (1) 277.5 μL of PBS, (2) 1.5 μL of 20 mM TBTA in DMSO, (3) 3 μL of 100 mM CuSO4 in diH_2_O, (4) 18 μL of 50 μM alkyne-AZDye 647 (Click Chemistry Tools #CCT-1301) in DMSO or 6 μL of 500 μM alkyne-AZDye 555 (Click Chemistry Tools #CCT-1289) in DMSO, and (5) 6 μL of 50 mM TCEP in diH2O. Once combined, each section was incubated in 50 μL of the reaction mix, protected from light, and rocked at 50 rpm for 1 hr at room temperature. The sections were then subjected to alternating 5 min washes with PBS (3x) and solution D (3x) at room temperature, followed by an additional set of washes (1 x 5 min and then 1 x 20 min) in solution D containing 10% DMSO and 50 mM EDTA. After additional (3 x 5 min) washes with PBS, the sections were either subjected to immunohistochemistry for α-actinin as described in the “Transverse Splitting” methods or stained for total protein with a No-Stain kit (Invitrogen #A44449). Specifically, 1 µL of the No-Stain Activator was added to the 500 µL of 1× No-Stain Labeling Buffer and vortexed. Then, 1 µL of the No-Stain Derivatizer was added to the solution and vortexed. Labeling was then performed by adding 50 µL of the prepared solution to each section, which was then incubated for 30 min at room temperature with rocking at 50 rpm. Labeled sections were then washed three times for 5 min with diH_2_O. In all cases, the washed sections were mounted with Prolong Diamond (ThermoFisher) and a 12 mm coverslip and then allowed to cure for 48 hr before imaging. Randomly selected 63x fields from the longitudinal sections were imaged with a Leica THUNDER Imager Tissue 3D microscope and subjected to deconvolution as described in the “Transverse Splitting” methods. For whole muscle cross-sections, images were collected with a BZ-X700 Keyence or a Leica THUNDER Imager Tissue 3D microscope. ImageJ was then used to assess the area of the entire muscle cross-section, as well as the intensity of the signal for the ANL-labeled proteins and total protein within each section. In some instances, the total number of muscle fibers per cross-section, along with the mean CSA of 70-120 randomly selected fibers per muscle, was also assessed as previously described (*31*). Notably, all image acquisitions and analyses were performed by investigators who were blinded to the experimental conditions of the samples.

### Determination of sarcomere diameter

Cross-sections (5 µm thick) from the central portion of immersion fixed and cryoprotected PLT muscles were collected on microscope slides with a cryostat that was chilled to -35°C. Immediately upon collection, the sections were transferred to diH_2_O water for 15 min to hydrate the samples. Residual water around the fully hydrated section was removed, and a hydrophobic circle was drawn around each section using a PAP pen (Thermo Fisher Scientific). Slides were then placed into a humidified box for all subsequent washing and incubation steps, which were all performed at room temperature on a rotating rocker set to 50 RPM. Sections were washed with PBS for 5 min and then incubated for 30 min in solution D and then overnight in solution D containing mouse IgG1 anti-SERCA1 (1:100, VE121G9, Santa Cruz #SC-58287) and rabbit anti-dystrophin (1:100, ThermoFisher #PA1-21011). The next day, sections were washed three times for 10 min and then three times for 30 min with PBS and subsequently incubated overnight in solution D containing AlexaFluor® 594 conjugated goat anti-mouse IgG, Fcγ Subclass 1 specific (1:2000, Jackson Immunoresearch #3115-585-205) and AlexaFluor® 488 conjugated goat anti-rabbit IgG (1:5000, Invitrogen #A11008). Following the overnight incubation, sections were washed three times for 10 min and then three times for 30 min with PBS. Washed sections were mounted with ProLong™ Gold Antifade Mountant (ThermoFisher) and covered with a 12 mm coverslip, and then allowed to cure for 24 hr before imaging. Our previously detailed Fluorescence Imaging of Myofibrils with Image Deconvolution (FIM-ID) workflow was then used to determine the minimal and maximal Feret diameter of the sarcomeres in 30 randomly selected fibers per muscle (*59*).

### Newly synthesized protein hot spot quantification

Longitudinal sections were subjected to a copper-catalyzed ‘click’ reaction with alkyne-AZDye 555 and immunohistochemistry for α-actinin, as detailed above. Randomly selected 63x fields were imaged with a Leica THUNDER Imager Tissue 3D microscope and subjected to deconvolution as described in the “Transverse Splitting” methods. The resulting images were then subjected to the various analyses that are detailed in Supplementary Material 2.

### Statistical analysis

Statistical significance was determined using student’s t-tests, paired t-tests, one-way ANOVA, or one-way repeated measures (RM) ANOVA with Tukey’s post hoc comparisons, two-way ANOVA, or two-way RM ANOVA with Fisher’s LSD post hoc comparisons. Samples that were more than three standard deviations from the mean within a given group were excluded as outliers. Differences between groups were considered significant when P < 0.05. All statistical analyses were performed with GraphPad Prism version 10.4.2 for Windows.

## Supporting information

Genotyping Primers

NSP Hot Spot Quantification Procedures

Uncropped Western Blot and Gels

## Funding

National Institutes of Health grant R01AR082816 (TAH)

National Institutes of Health grant R01AR074932 (TAH)

National Institutes of Health grant R56AR057347 (TAH)

## Author contributions

Conceptualization: JEH, KWJ, HGP, TAH

Methodology: JEH, KWJ, HGP, MM, CGKF, TAH

Investigation: JEH, KWJ, RKAS, HGP, MM, CGKF, ANL, GTL, PMF, TAH

Formal Analysis: JEH, KWJ, RKAS, HGP, MM, CGKF, GTL, TAH

Visualization: JEH, KWJ, HGP, GTL, TAH

Supervision: TAH

Writing - original draft: JEH, KWJ, TAH

Writing - review & editing: JEH, KWJ, RKAS, HGP, MM, CGKF, ANL, GTL, PMF, TAH

## Competing interests

Authors declare that they have no competing interests.

## Data Availability

All data are available in the main text or supplementary material files.

## Electronic Supplementary Material

Supplementary Material 1 – Genotyping Primers

Supplementary Material 2 – NSP Hot Spot Quantification Procedures

Supplementary Material 3 – Supplemental Figures

Supplementary Material 4 – Supplemental Video 1

Supplementary Material 5 – Supplemental Video 2

Supplementary Material 6 – Uncropped Western Blot and Gel Images

Supplementary Material 7 – Source Data

## Notes

### Competing Interest Statement

The authors have declared no competing interest.

### Summary of Updates

Minor changes to the discussion were made following feedback that was received from the public after the initial release

