## Supplementary material for "Mechanical Loading Induces the Longitudinal Growth of Muscle Fibers via an mTORC1-Independent Mechanism": Genotyping Primers

**Supplemental Table 1**

| Target | Sequence (5' → 3') |
|---|---|
| CMV-CRE Forward Primer (JAX oIMR1084) | GCG GTC TGG CAG TAA AAA CTA TC |
| CMV-CRE Reverse Primer (JAX oIMR1085) | GTG AAA CAG CAT TGC TGT CAC TT |
| Wild-type (intact Rosa26) Forward Primer (JAX 26209) | CTG GCT TCT GAG GAC CG |
| Wild-type (intact Rosa26) Reverse Primer (JAX oIMR9021) | CCG AAA ATC TGT GGG AAG TC |
| $MetRS^{L274G}$ Mutant Forward Primer (JAX 12614) | ACC ACT ACC AGC AGA ACA CC |
| $MetRS^{L274G}$ Mutant Reverse Primer (JAX 26209) | GGC AGA TTG CAC TAG CAG AG |
