## Supplementary material for "Mechanical Loading Induces the Longitudinal Growth of Muscle Fibers via an mTORC1-Independent Mechanism": NSP Hot Spot Quantification Procedures

### <u>Supplementary Material 2</u>: Newly Synthesized Protein (NSP) Hot Spot Quantification Procedures

Due to the subjective nature of these analyses, the investigators acquiring the images, as well as those who performed the analyses, were blinded to the experimental condition of the samples.

#### <u>NSP Hot Spot Positive Fibers</u>

Fibers were categorized as positive for NSP hot spots if they contained multiple loci that were densely populated with NSPs. To illustrate this, four 63x images of NSPs in muscles that were subjected to mechanical overload (MOV) or the sham condition are shown below. Asterisks indicate fibers that were categorized as positive for NSP hot spots.

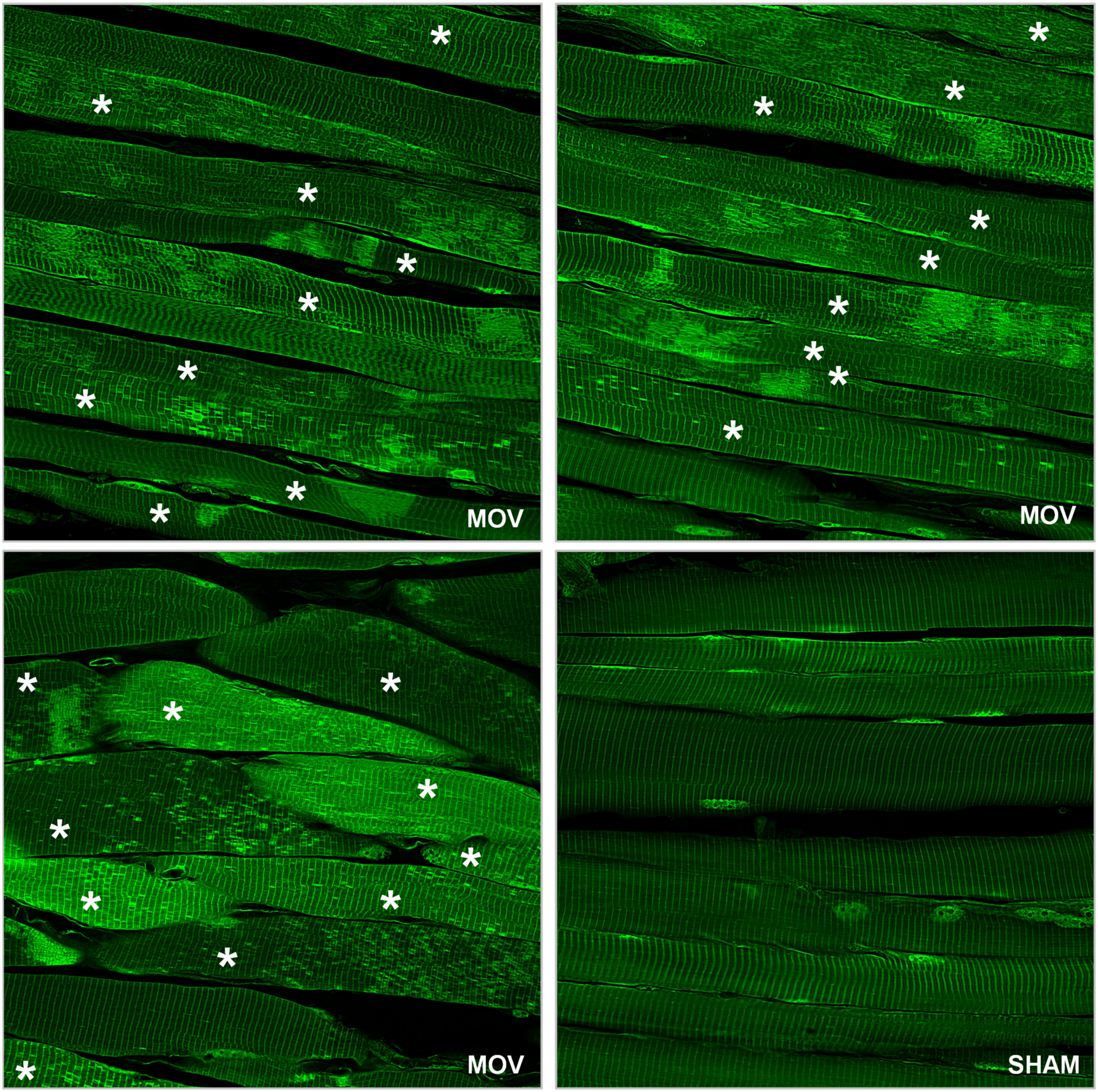

### Normal vs. Disarrayed ROIs

Fibers that were categorized as positive for NSP hot spots were screened for the presence of “normal” and “disarrayed” 100 x 100 pixel ROIs (100 $\mu m^2$) when viewing the signal for α-actinin. ROIs that were classified as “disarrayed” contained highly discontinuous Z-lines. When selecting these ROIs, preference was given to regions with the highest discontinuity and/or the appearance of numerous “Y” shaped splits. The image below illustrates the priority ranking of four different “disarrayed” ROIs with a ranking of 1 indicating the top preference.

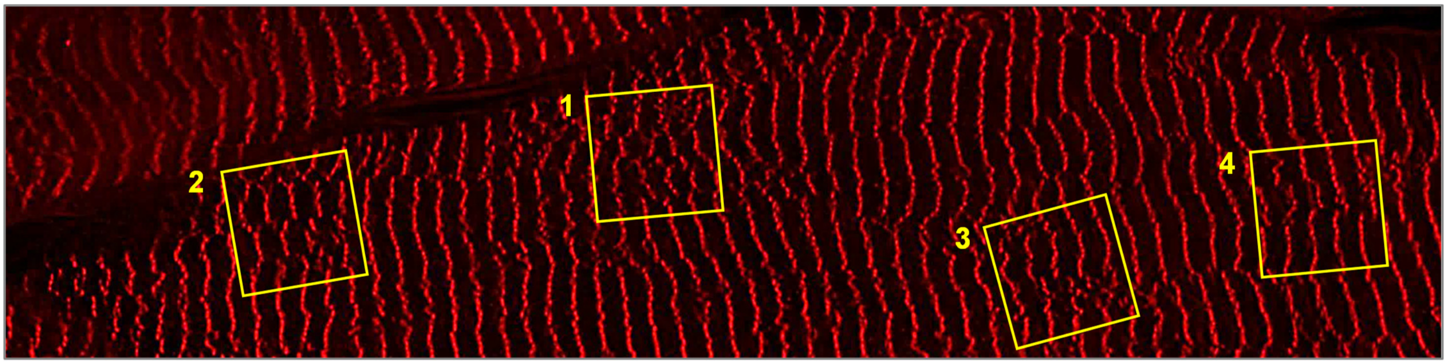


ROIs that were classified as “normal” contained continuous Z-lines. When selecting these ROIs, efforts were made to maximize the distance of the ROI from surrounding areas of disarray. For example, the image below shows three “normal” ROIs (white boxes labeled A-C), as well as three disarrayed ROIs (yellow boxes labeled D-F). In this example, the top-ranking “normal” ROI is A, followed by C, and then B.

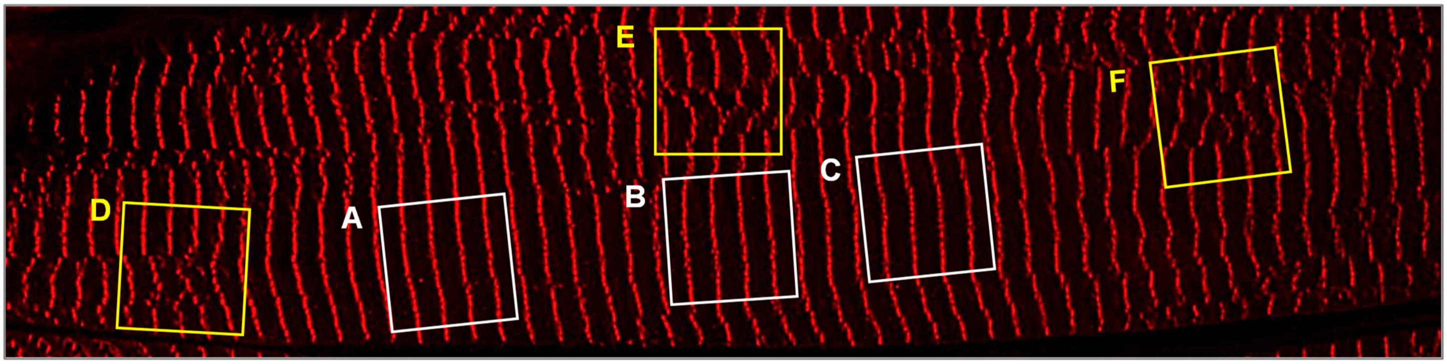


Fibers that contained both “normal” and “disarrayed” ROIs were subjected to further quantitative analyses. Specifically, an equivalent number of “normal” and “disarrayed” ROIs within each fiber were identified (up to 3 of each type of ROI per fiber). Then, within each ROI, the periphery of the individual hot spots was manually traced and the total number of hot spots per ROI was recorded. As illustrated on the next page, some fibers were permissive to the analysis of multiple ROIs and, in these instances, the mean values for the “normal” and “disarrayed” ROIs within the fiber were calculated. In all cases, the final “normal” and “disarrayed” ROI data for each fiber were recorded as paired values. Notably, several of the fibers that were categorized as positive for NSP hot spots contained an extensive number of “disarrayed” ROIs but did not contain a “normal” ROI for a paired analysis, and for this reason, ROI level quantification on these fibers was not performed.

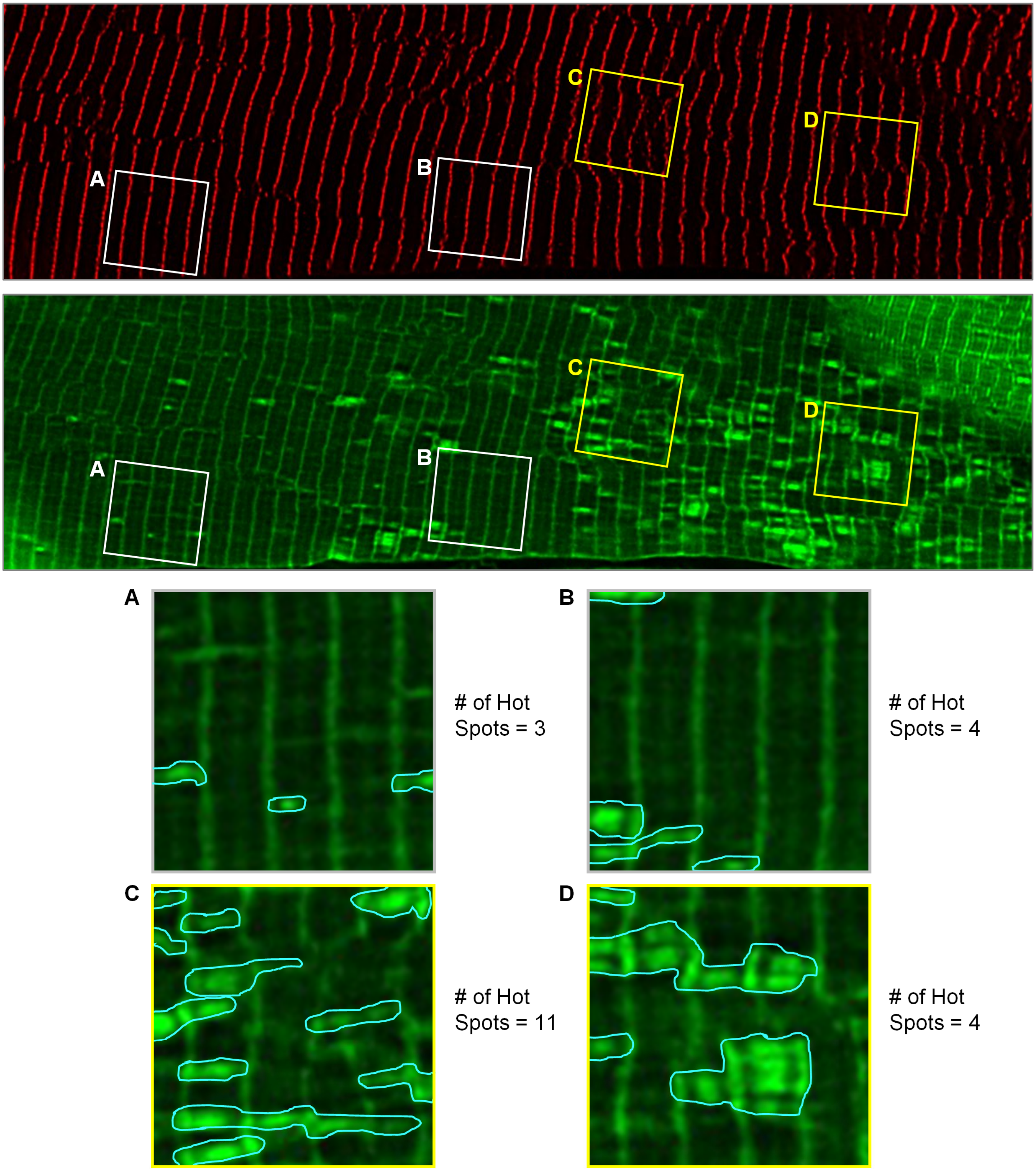


Paired Values Reported for the Fiber:
Normal ROI: # of Hot Spots = 3.5, Disarrayed ROI: # of Hot Spots = 7.5

### Classification of NSP Hot Spot Morphology

To be included in the classification analysis, an NSP hot spot needed to have a signal intensity at least 2-fold higher than the local background and a diameter between 400 and 1900 nm (corresponding to the values that represents the 0.5$^{th}$ percentile of the minimal and 99.5$^{th}$ percentile of the maximal Feret diameters in sarcomeres of plantaris muscles that had been subjected to 8 days of MOV; see Supplemental Figure 1). All fibers within a given 63x field that contained at least 15 qualifying NSP hot spots were analyzed. Within each of these fibers, at least 15 and up to 30 qualifying NSP hot spots were randomly selected and classified according to whether their morphology conformed to one of three previously proposed models of in-series sarcomerogenesis (*1-3*).

The first model we considered is based on a transverse Z-line splitting event that was recently described by Rodier et al. (2025) (*1*). Specifically, this model proposes that in-series sarcomerogenesis is initiated when titin / thick filament complexes detach from the Z-lines of an existing sarcomere. Equal portions of these complexes are then pulled in opposite directions, and lead to the exposure of binding sites that facilitate the recruitment of new sarcomeric proteins. As illustrated in Figure 7A, if the recruited proteins consist of NSPs, then this process would give rise to exactly two in-series sarcomeres that are densely populated with NSPs. An example of an NSP hot spot (green) exhibiting this morphology is shown within the "A" bracket in the adjacent image. All NSP hot spots that displayed this morphology were classified as <u>Rodier et al. Z-line splits</u>.

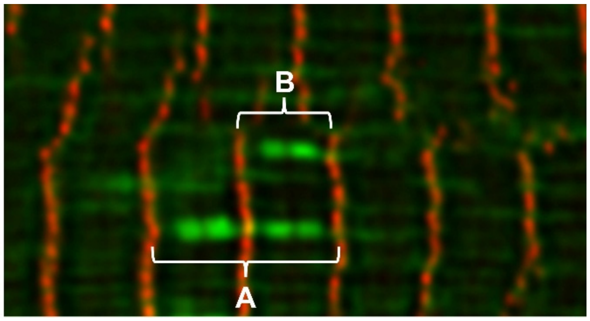


The second model of in-series sarcomerogenesis that we considered is based on a transverse Z-line splitting event originally described by Yu et al. (2003) (*2*). Here, the process is proposed to begin with the breakdown and broadening of a single Z-line. As this process continues, the remaining components of the Z-line get pulled in opposite directions, which allows new sarcomeric proteins to be incorporated into the region of expansion. As illustrated in Figure 7B, if these sarcomeric proteins are NSPs, then this process would result in the formation of a single sarcomere that is densely populated with NSPs. An example of an NSP hot spot whose morphology is consistent with this model is shown within the "B" bracket above, and all such NSP hot spots were classified as <u>Yu et al. Z-line splits</u>.

The final model of in-series sarcomerogenesis we considered was proposed by Jahromi and Charlton 1979 (*3*) and involves transverse splitting at the H-zone. More precisely, the model argues that thick filaments are bisected at the H-zone, after which the two halves get pulled in opposite directions. As this occurs, it is thought that new myosin molecules get incorporated at the severed ends of the thick filaments while new thin filament / Z-line complexes form at the site that was previously occupied by the M-line. Importantly, if the incorporated proteins are NSPs, then it will result in the formation of an NSP hot spot that is confined to the adjacent inner halves in a pair of in-series sarcomeres (Fig. 7C). An example of such an NSP hot spot is shown within the "C" bracket in the adjacent image, and all hot spots that displayed this type of morphology were classified as <u>H-zone splits.</u>

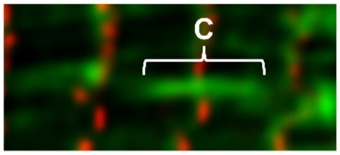


During the development of our classification procedure, it became evident that a subset of the qualifying NSP hot spots did not align with any of the previously described models of in-series sarcomerogenesis. As such, we created two additional classes: <u>Short – Atypical</u> and <u>Long – Atypical</u>. Short – Atypical hot spots were defined as being less than two in-series sarcomeres in length, and, as illustrated in the examples below, they typically spanned either ~1/2 sarcomere "D" or ~1.5 sarcomeres "E". In contrast, Long – Atypical hot spots were defined as exceeding two in-series sarcomeres in length and, as shown under "F", they were typically composed of ≥3 in-series sarcomeres.

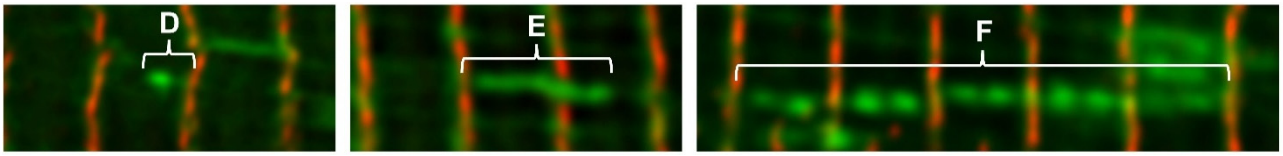


After 15 - 30 qualifying NSP hot spots per fiber had been classified, the percentage within each class was calculated, and the per-fiber results were reported in the final datasets.
