## Supplemental Figures for "Mechanical Loading Induces the Longitudinal Growth of Muscle Fibers via an mTORC1-Independent Mechanism"

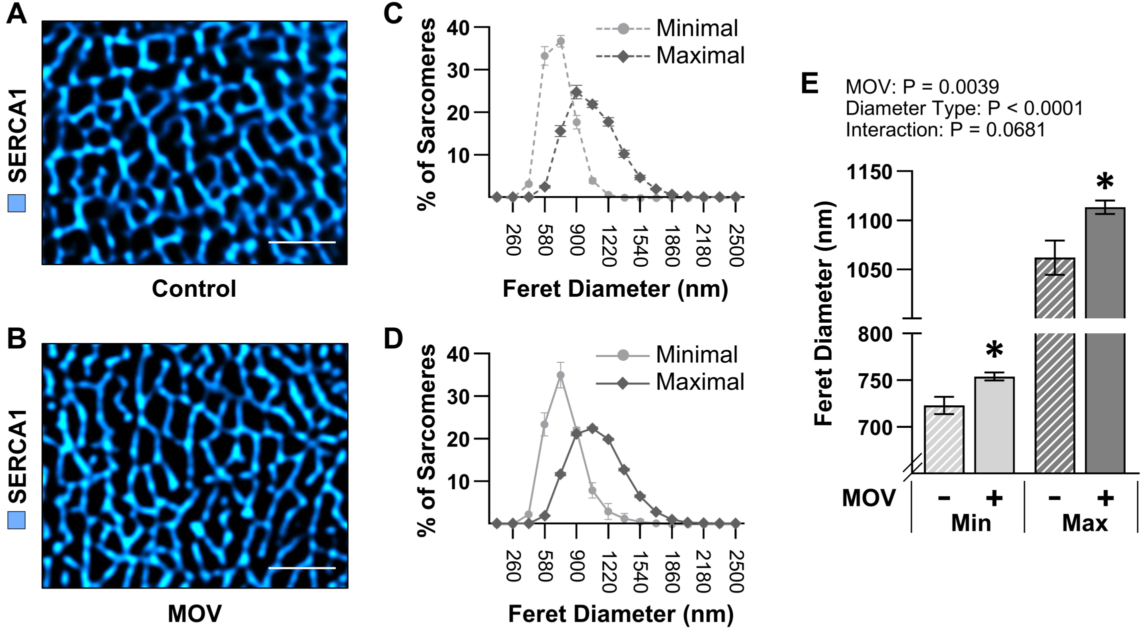


**Supplemental Figure 1. The Effect of MOV on Sarcomere Diameter.** C57BL/6J mice were subjected to a mechanical overload (MOV) or sham (control) surgery. (**A**-**B**) After 8 days, the plantaris muscles were collected and then mid-belly cross-sections were subjected to immunohistochemistry for dystrophin to identify the periphery of the muscle fibers (not shown) and SERCA1 to identify the periphery of the sarcomeres (cyan). The minimal and maximal Feret diameters of the sarcomeres within randomly selected fibers was determined as detailed in the methods section. Scale bar = 2 µm. (**C**-**D**) Frequency distribution of the minimal (light gray) and maximal (dark gray) Feret diameters of the sarcomeres in the control (**C**) and MOV (**D**) muscles. For the control condition, n = 37868 sarcomeres from 4 independent muscles (5330-16329 sarcomeres per muscle). For the MOV condition, n = 100360 sarcomeres from 9 independent muscles (5687-14103 sarcomeres per muscle). (**E**) The mean values for each of the muscles in C-D were analyzed with two-way RM ANOVA. * Significantly different from control within a given Feret type, $P < 0.05$.

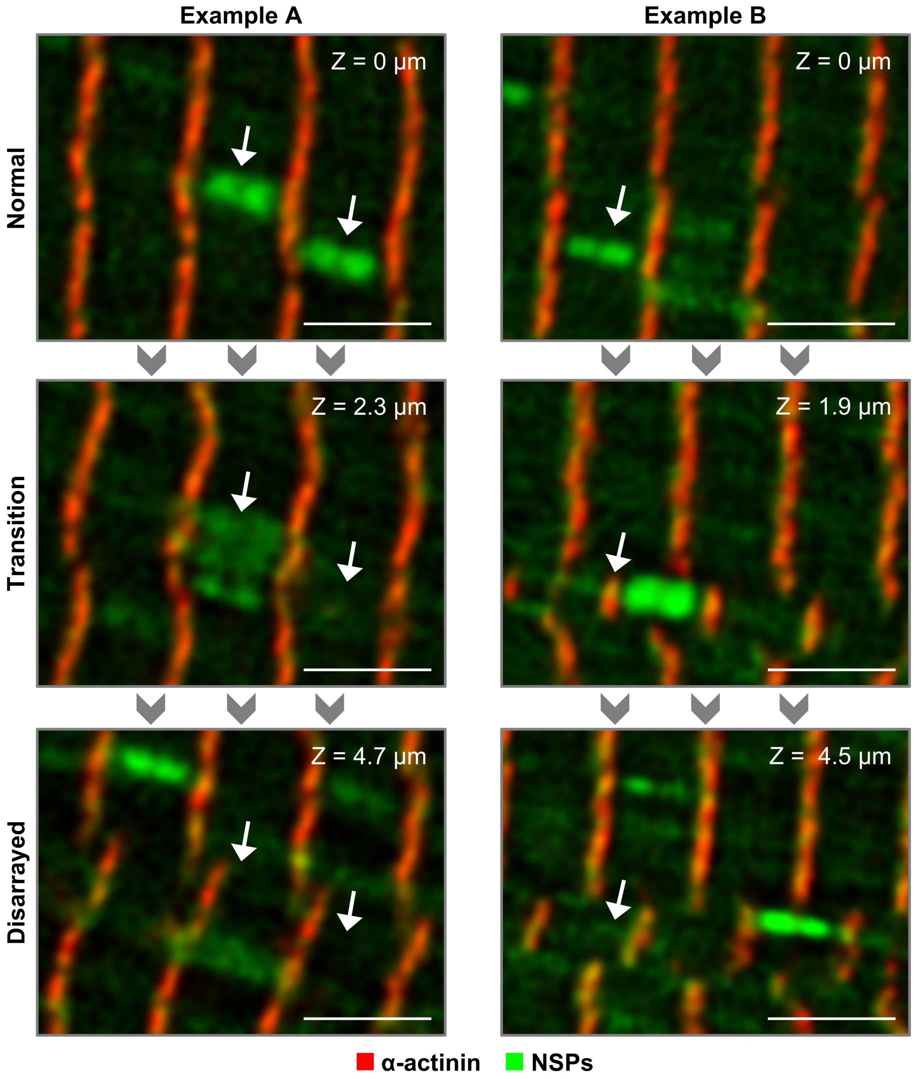


**Supplemental Figure 2. Three-dimensional Assessment of NSP Hot Spots.** MetRS$^{L274G+/+}$ mice were subjected to a mechanical overload (MOV) surgery, and after 7 days, the mice were injected with ANL. The plantaris muscles were collected 24 hr later, and then thick (10 µm) longitudinal sections were subjected to immunohistochemistry for α-actinin (red) and a click reaction with alkyne-AZDye 555 (green) to label the newly synthesized proteins (NSPs). A representative region of interest was identified and then imaged at the indicated depths of the Z-plane. Arrows point to the same location along the X and Y planes. Two examples are provided to illustrate how NSP hot spots in a region with a "normal" Z-line configuration reside immediately superficial to a region of "disarray". Scale bars = 3 µm.

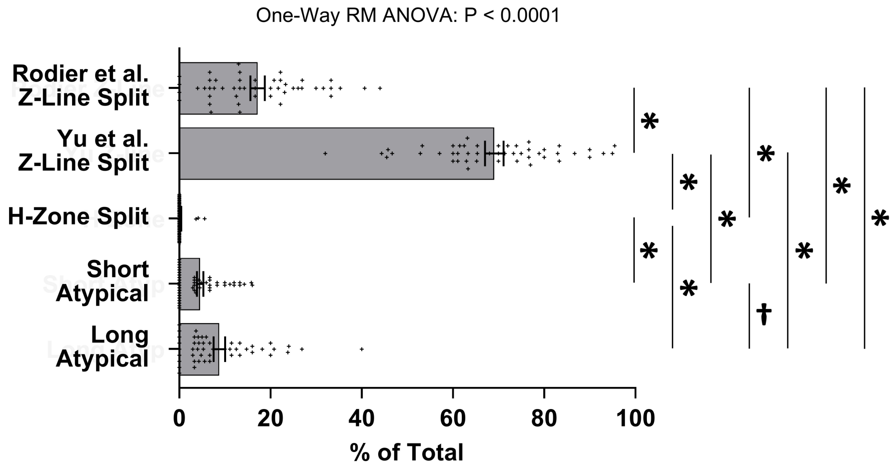


**Supplemental Figure 3. Distribution of sarcomere-level NSP hot spot morphologies in muscles subjected to mechanical overload.** $MetRS^{L274G+/+}$ mice were subjected to a mechanical overload (MOV), and after 7 days, the mice were injected with ANL. The plantaris muscles were collected 24 hr later and longitudinal sections were subjected to immunohistochemistry for α-actinin and a click reaction with alkyne-AZDye 555 to label the NSPs. The images were assessed for the presence of fibers that had qualifying NSP hot spots (e.g., signal intensity at least 2-fold higher than the local background and dimensions that were consistent those of a sarcomere) and, for each analyzed fiber, the proportion of the NSP hot spots whose morphology aligned with the models of in-series sarcomerogenesis or one of the two atypical split types was determined, n = 48 fibers from 15-30 NSP hot spots / fiber, 10-15 fibers / muscle, 3 muscles (see Supplemental Material 2 for details). Data are presented as individual fiber values as well as group means ± SEM. The data were analyzed with one-way RM ANOVA. * Significant difference between the indicated groups, † $P < 0.05$, * $P < 0.001$.

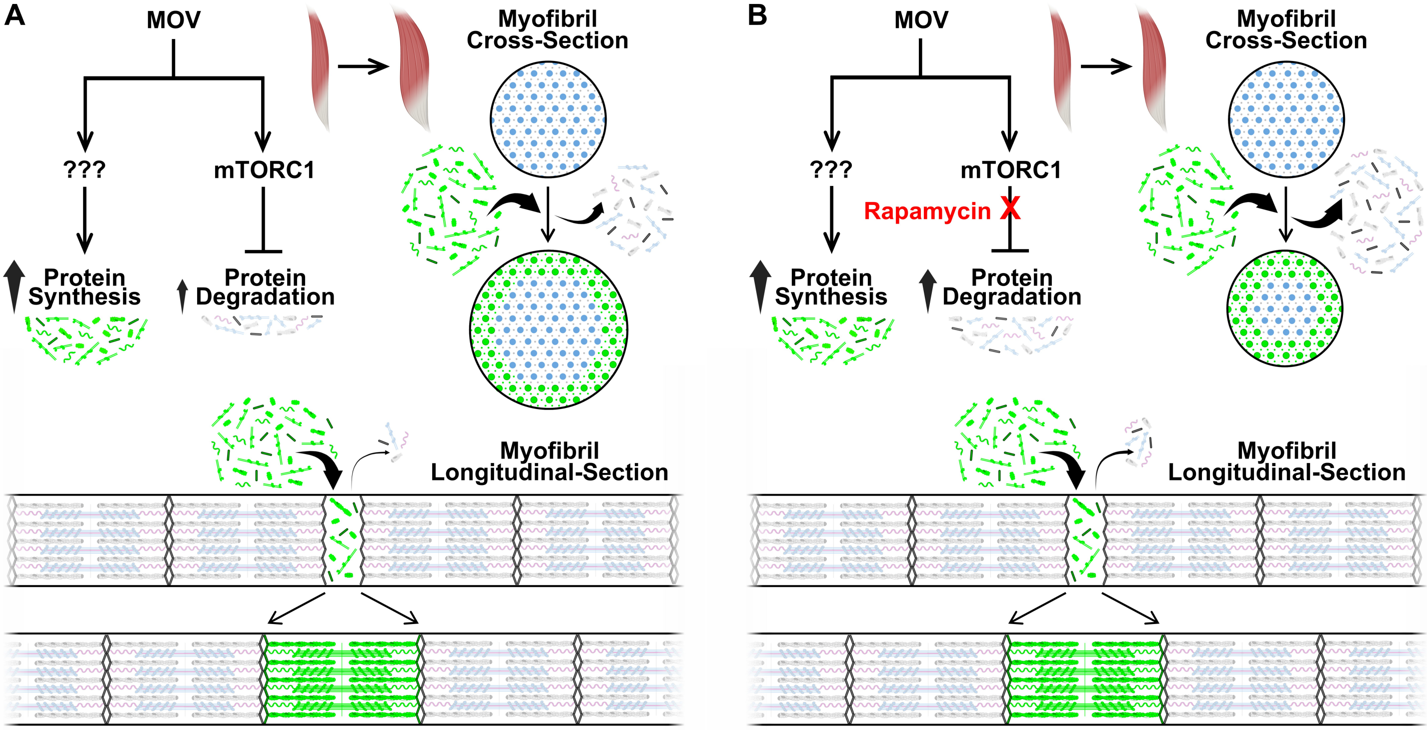


**Supplemental Figure 4. Illustration of How the Inhibition of mTORC1 Could Differentially Regulate Radial and Longitudinal Growth.** The hypothetical model shows plantaris muscles along with cross-sectional and longitudinal views of myofibrils that are composed of pre-existing proteins such as Z-lines (dark gray), titin (light pink), thick filaments (light blue), and thin filaments (gray), as well as newly synthesized proteins (green). (**A**) The hypothetical model predicts that under normal conditions, mechanical overload (MOV) induces a robust increase in protein synthesis through a currently unknown mechanism. MOV also induces robust activation of mTORC1, and this exerts a suppressive effect on protein degradation. As a result, the rate of protein synthesis greatly exceeds the rate of protein degradation, and this leads to a large increase in muscle mass. At the ultrastructural level, the increase in muscle mass can be attributed to an increase in both the diameter (i.e., radial growth) and length (i.e., longitudinal growth) of the myofibrils. Importantly, in the model, it is assumed that the radial growth results from a moderate net increase in the synthesis of new proteins relative to the degradation of the pre-existing proteins. On the other hand, the longitudinal growth is mediated by the in-series addition of sarcomeres, which is almost entirely driven by the synthesis of new proteins (i.e., very few pre-existing proteins are degraded during this process). (**B**) When signaling through mTORC1 is inhibited (e.g., by rapamycin), the MOV-induced increase in protein synthesis is unaffected, but the suppressive effect of mTORC1 on protein degradation is lost. As a result, the overall rate of protein synthesis only moderately exceeds the rate of protein degradation, and thus, the resulting increase in muscle mass is reduced when compared with what occurs in the normal condition. At the ultrastructural level, the smaller increase in muscle mass can be attributed to the loss of radial growth. Specifically, the enhanced degradation of the pre-existing proteins offsets the accumulation of the newly synthesized proteins and thus, the diameter of the myofibrils does not change. Conversely, the degradation of pre-existing proteins does not make a major contribution to the process via which new in-series sarcomeres are added. As such, the increase in protein degradation would exert little, if any, effect on the induction of longitudinal growth.
