## Supplementary material for "Mechanical Loading Induces the Longitudinal Growth of Muscle Fibers via an mTORC1-Independent Mechanism": Uncropped Western Blot and Gels

### Uncropped Western Blot and Gel Images – Figure 4

**Figure 4B**

Anti-GFP

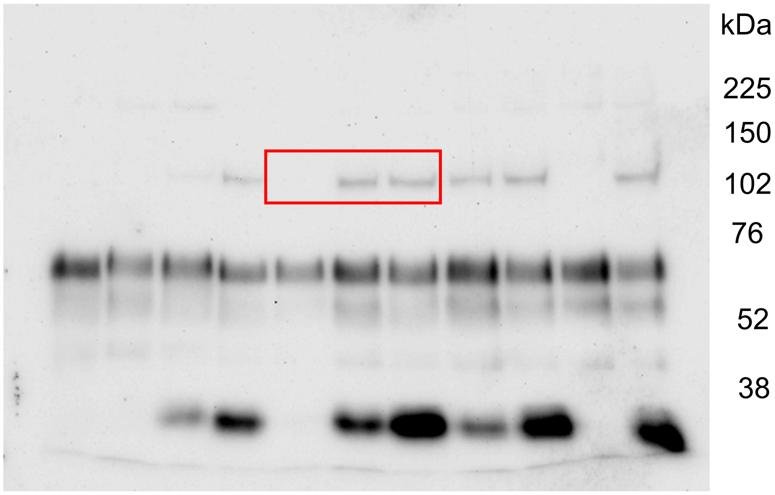


Coomassie Blue = Total Protein

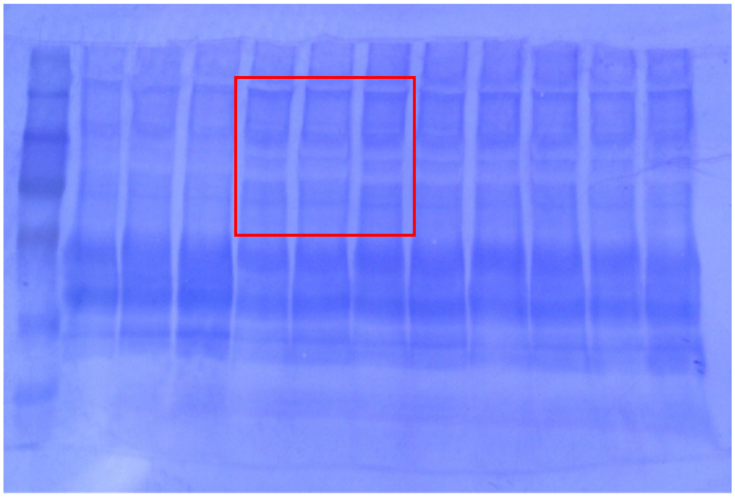

**Figure 4C**

DBCO-Cy5.5

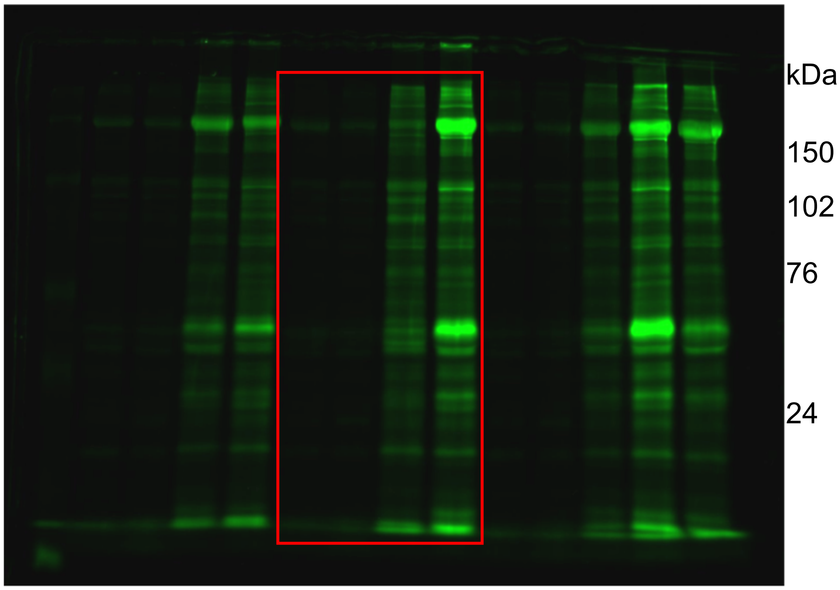


No Stain = Total Protein

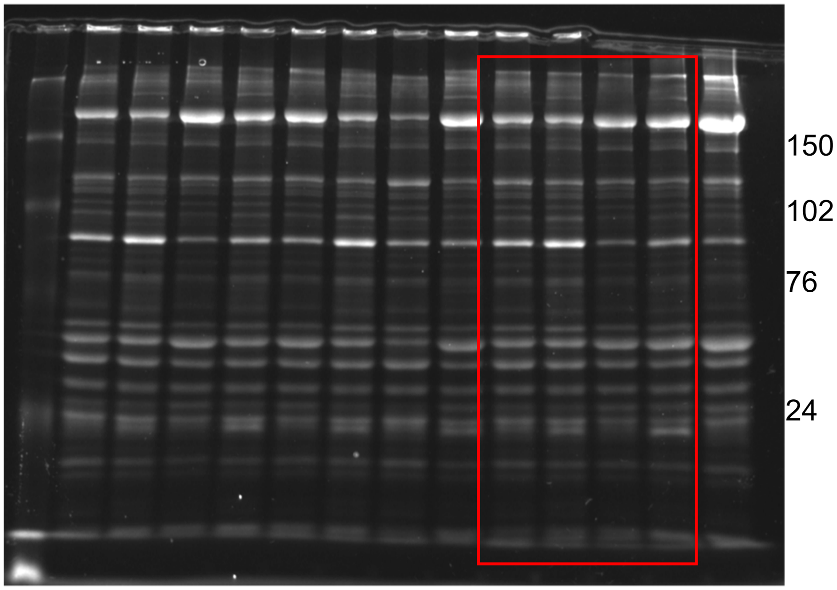

### Uncropped Western Blot and Gel Images – Figure 6

**Figure 6E**

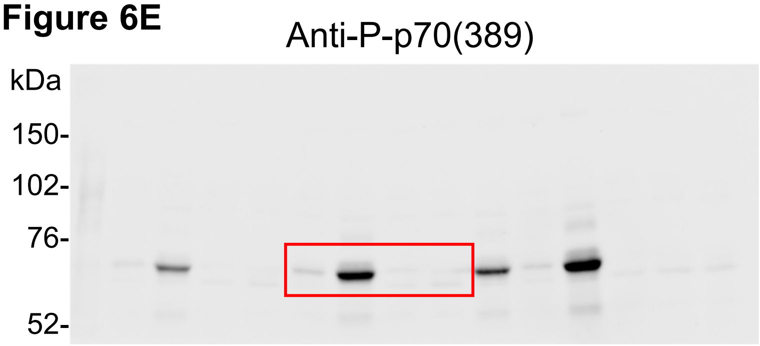


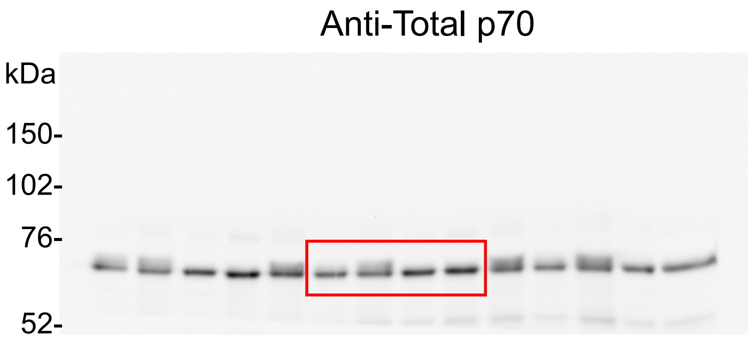


**Figure 6F**

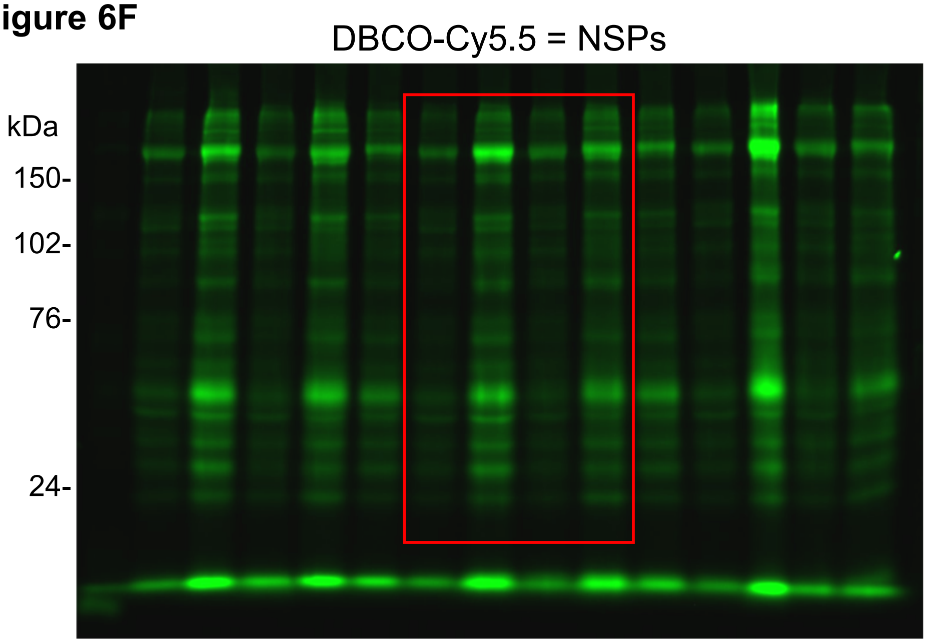


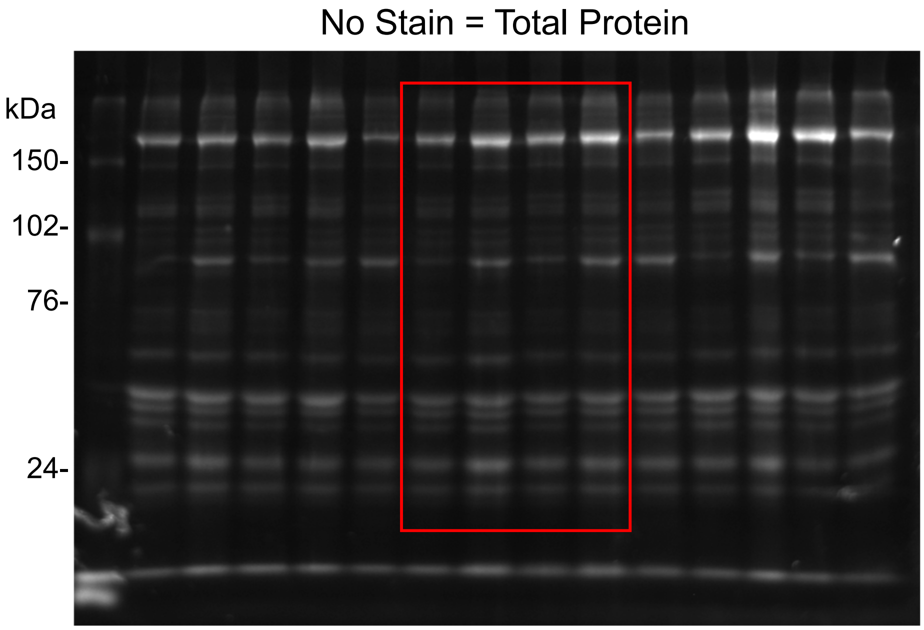
